# Laser patterning bioprinting using a light sheet-based system equipped with light sheet imaging produces long-term viable skin constructs

**DOI:** 10.1101/2023.05.10.539793

**Authors:** Levin Hafa, Louise Breideband, Lucas Ramirez Posada, Núria Torras, Elena Martinez, Ernst H.K. Stelzer, Francesco Pampaloni

**Author notes:** Equal contribution (Levin Hafa, Louise Breideband).

## Abstract

This research introduces a new 3D bioprinter that incorporates live imaging of the bioprinted tissue with high resolution and high-speed capabilities. The printer employs a light sheet-based system to photocrosslink polymers into hydrogels at a printing speed of up to 0.66 mm³/s with a resolution of 15.7 µm. A significant advancement of this bioprinter is its ability to track cells and bioink during crosslinking, which enables real-time evaluation of the 3D-bioprinted structure’s quality. Fibroblast cells were encapsulated using this method, and the viability was evaluated directly after bioprinting and seven days after encapsulation, which was found to be high (83% ± 4.34%). Furthermore, a full-thickness skin construct was bioprinted and maintained in culture for 6 weeks, demonstrating the long-term viability and physiological relevance of the bioprinted tissue. The usage of solid-state laser beam scanning devices could enhance bioprinting’s speed and precision. This fast and accurate light-based bioprinter offers a promising platform for generating customizable 3D-printed structures with viable long-term cultures.

**Teaser:** A novel bioprinter with live imaging capability using light sheet microscopy produces viable long-term cultures with high-resolution structures.

**Graphical abstract:** General workflow of bioprinting skin constructs using light sheet bioprinting.

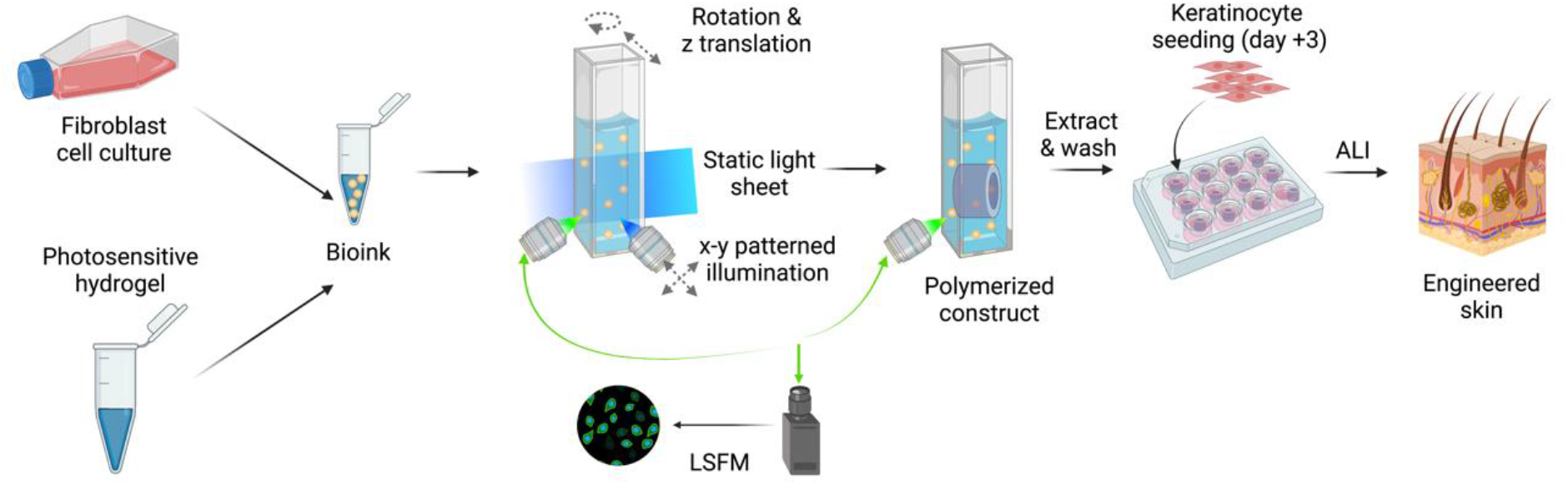

## Introduction

The field of tissue engineering is a rapidly developing interdisciplinary area that offers substantial potential. Advancements in techniques, materials, and culture methods are being made continuously, and the expectations for tissue engineering products are high. Such products hold the promise of replacing animal models for basic research and drug discovery, as well as facilitating tissue regeneration and organ transplantation. Animal models, despite being essential in research, are deficient in accurately representing human physiology and molecular processes (*1,2*). Ethical concerns and increasingly stringent regulations promote the replacement of animal models when the principles of reduction and refinement do not apply (the 3R concept) (*3*). The recent enactment of the U.S. Food and Drug Administration (FDA) Modernization Act 2.0, which authorizes the use of alternatives to animal testing in the drug discovery process, underscores the importance of tissue engineering in the pharmaceutical industry (https://www.congress.gov/bill/117th-congress/senate-bill/5002 (*4*)). Although human donors are the primary source of organs for transplant (allotransplantation), only 20% of individuals registered on the US National Transplant Waiting List received a transplant in 2020, despite advances in transplantation techniques (*5*). Xenotransplantation, particularly from pigs, has been investigated as an alternative source for organ production. Nevertheless, xenotransplantation poses substantial challenges, such as the potential for infectious complications and extensive preventative and curative treatment regimens for patients (*6*).

Among the biofabrication techniques, three-dimensional (3D) bioprinting offers design flexibility, reproducibility, and high level of detail (*7*). First developed for practical purposes by Thomas Boland’s group in 2003, the system was defined as “computer-aided, jet-based 3D tissue-engineering of living human organs” (*8*). This technique was developed as a faster, more accurate alternative to classic tissue engineering technologies (for example, 3D cell culture in drops of an extracellular matrix like Matrigel) (*9*). Since its inception, 3D bioprinting has evolved and branched into several categories, in which 3D organization is achieved by different techniques. The branch of bioprinting that achieves material deposition using physical pressure through a nozzle is divided into extrusion and inkjet bioprinting (*10, 11*). The former deposits a constant line of material while the latter deposits droplets of biomaterial. The speed (60 mm/s for extrusion (*12*)) and resolution of nozzle-based bioprinting depends on the velocity and diameter of the nozzle, respectively. Those methods are limited by the shear pressure imposed by the nozzle which reduces the possible range of cell density and material viscosity (*12*).

Another branch of bioprinting utilizes light to produce objects (a process called photocrosslinking) (*13*). Digital light projection (DLP) uses light projected onto a platform to crosslink entire planes at once. These planes can also be generated by radon transform to provide reverse-computerized tomography (CT) stacks which are projected into a volume of photocrosslinkable polymer, a principle on which volumetric bioprinting is based (*14*). The former method enables fast bioprinting (in the order of a mm³/s) (*15, 16*) with good resolution (30 µm to 50 µm) (*16, 17*) and is not limited by the viscosity of the polymer (*12*). Xolography is another volumetric 3D printing method worth noting, which has similar optical characteristics to this publication. There, a projector shines a 2D image into a resin-filled cuvette and two orthogonal static light sheets activate the photoinitiator in the plane being crosslinked. By superpositioning the light sheets with the projections, the resolution in the Z-plane can be increased (*18*). The highest resolution can be achieved with a two-photon light source as a trigger for the photocrosslinking, reaching a resolution of 0.1 µm (*19*). Higher speeds of maximum 20 mm/s can be achieved with this method, although the resolution in this case is around 250 µm (*20*).

While the field of bioprinting has been focusing on speed and resolution, the assessment of cell viability and function within the bioprinted tissue are done “offline” in a separate device. Therefore, bioprinting and imaging are currently two separate processes in most devices. Exceptions exist, some that combine live brightfield monitoring of the process (*16, 21, 22*). Nonetheless, they do not allow for online monitoring of both the hydrogel or the cells and, so far, no mention of an integrated fluorescent imaging device has been made.

In this work, we present a method that encompasses high printing speed (0.66 mm³/s) and high resolution (15.7 µm) while introducing a fully integrated and streamlined fluorescent light sheet microscope. Using the principle of direct laser patterning, a gaussian light beam is patterned at high velocity onto a vat of photocrosslinkable material. To control the z-resolution, a static light sheet is projected at a 90° angle to the patterned light beam, having a limited volume where the intensity contribution between the two light sources, after a pre-determined time, allows to surpass the dosage threshold needed to trigger the crosslinking process, thus allowing for a confined voxel to be crosslinked. The patterned light beam, in theory, allows for a 11 µm x- and y-resolution (FWHM of beam at focal point, data not shown) and a 49 µm z-resolution (FWHM of the light sheet, Fig. S2.). In practice, 15.7 µm-sized objects have been printed.

Table 1 compares key properties of the light sheet bioprinter with recent 3D (bio-) printers. An extensive comparison of all important properties in a 3D (bio-) printer can be found in Supplementary Table S8.

**Table 1:**
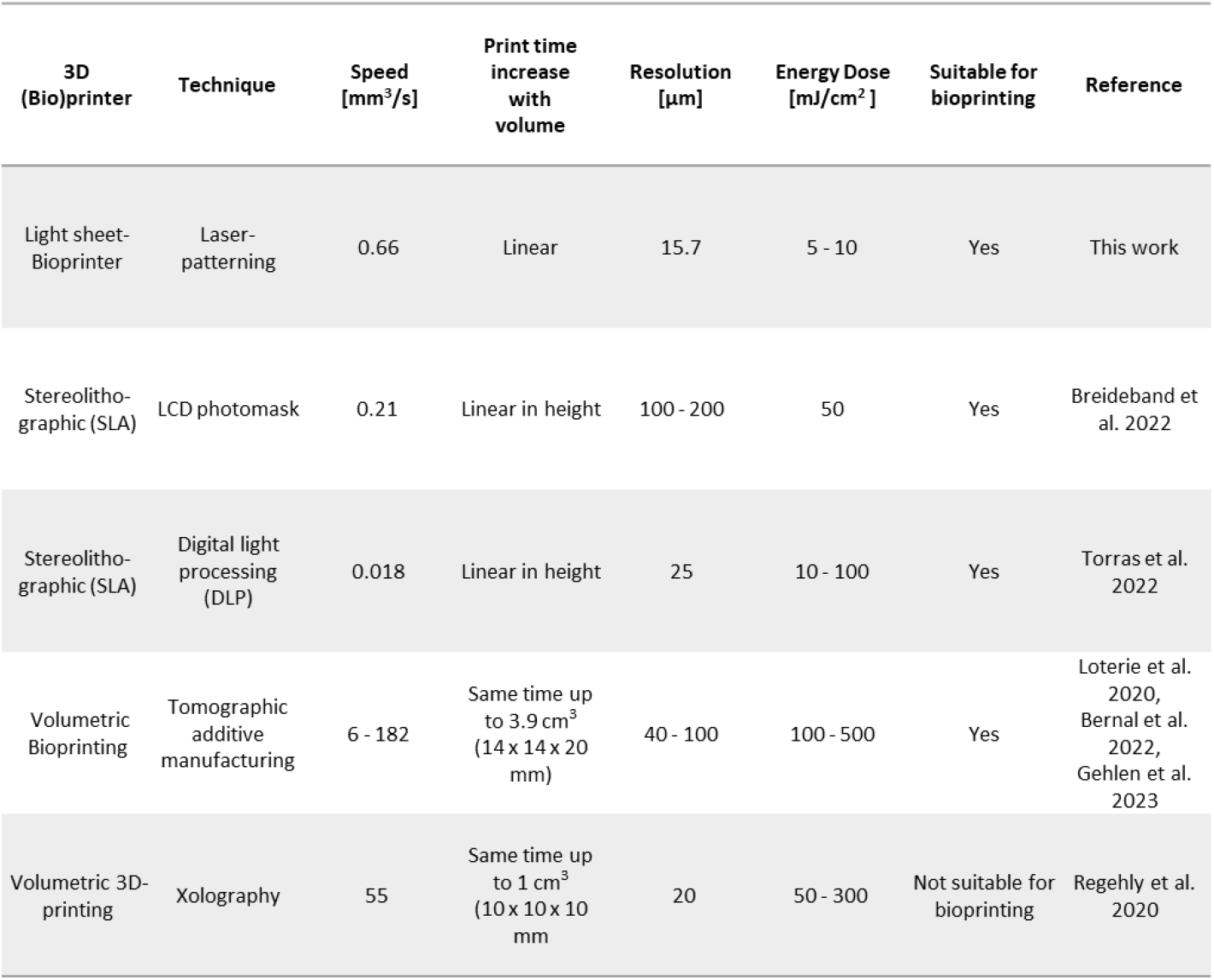
Comparison of 3D (Bio-)printers.

| 3D (Bio)printer | Technique | Speed [mm<sup>3</sup>/s] | Print time increase with volume | Resolution [μm] | Energy Dose [mJ/cm<sup>2</sup>] | Suitable for bioprinting | Reference |
| --- | --- | --- | --- | --- | --- | --- | --- |
| Light sheet-Bioprinter | Laser-patterning | 0.66 | Linear | 15.7 | 5 - 10 | Yes | This work |
| Stereolitho-graphic (SLA) | LCD photomask | 0.21 | Linear in height | 100 - 200 | 50 | Yes | Breideband et al. 2022 |
| Stereolitho-graphic (SLA) | Digital light processing (DLP) | 0.018 | Linear in height | 25 | 10 - 100 | Yes | Torras et al. 2022 |
| Volumetric Bioprinting | Tomographic additive manufacturing | 6 - 182 | Same time up to 3.9 cm<sup>3</sup> (14 x 14 x 20 mm) | 40 - 100 | 100 - 500 | Yes | Loterie et al. 2020, Bernal et al. 2022, Gehlen et al. 2023 |
| Volumetric 3D-printing | Xolography | 55 | Same time up to 1 cm<sup>3</sup> (10 x 10 x 10 mm) | 20 | 50 - 300 | Not suitable for bioprinting | Regehly et al. 2020 |

Additionally, the device integrates a light sheet microscope setting. This feature permits the observation of the crosslinking status of the hydrogel using fluorescent recovery after photobleaching (FRAP). Moreover, fluorescent cells before and after 3D bioprinting could be imaged. Human fibroblasts were encapsulated in a hydrogel based on thiol-ene click chemistry by bioprinting a hollow cylinder with visible light (ca. 8 mm³ printed within minutes). The process was fast and high resolution and produced a cell-laden construct that exhibited high short- and long-term cell viability, while conserving cell functionality, as is demonstrated by the presence of typical dermal markers. Full-thickness skin constructs (encapsulated fibroblasts and subsequent co-culture with human keratinocytes in air-liquid conditions) were still viable at 41 days post-bioprinting and displayed epidermal and dermal characteristics. We demonstrated that light sheet bioprinting is capable of high speed and definition, with capabilities for even higher velocity and resolution. Additionally, the successful imaging of cells and hydrogels in a streamlined fashion using the bioprinting device opens an array of opportunities for biologists. This work aims to pave the way for improvements in the field of light-based bioprinting. By combining advanced laser scanning devices, such as acousto-optic modulators (AOM) and deflectors (AOD) to such a system developed in this work, printing resolution and speed can be improved even further.

## Results

### Combining a light sheet microscope with a custom-made bioprinting device

Light sheet fluorescence microscopy (LSFM) was effectively developed in the early 2000s as a selective/single plane illumination microscope (SPIM) (*23, 24*), using a cylindrical lens to create a coherent static light sheet and achieving a 3D scan by moving the specimen either in z- or θ-axis (depth and rotation, respectively). Later, light sheet systems have evolved to more dynamic processes using a galvanometer mirror to vertically (y-axis) scan an incoherent laser beam, resulting in digital scanned light sheet microscopes (DSLM) (*25, 26*). The system developed in this work further exploits the patterning implemented in the DSLM (*26*), one for the scanning in x-axis and one for the y-axis, and the stage movement in the z-axis to create three-dimensional light-beam patterns. The patterned light (395 nm) was directed through a scan, a tube, and an objective lens before entering the water filled specimen chamber in which a specimen holder (or cuvette) containing a light-sensitive bioink (composed by a photocrosslinkable hydrogel with or without cells) was suspended (see Figure 1a). The photocrosslinkable hydrogel, under the right conditions (wavelength, laser intensity and exposure time surpassing the crosslinking threshold of the hydrogel), crosslinked, resulting in a bioprinted object either free-floating in the non-crosslinked material or attached to a support. In addition to the bioprinting application, the galvanometer mirror, if scanned only in the y-axis, resulted in a conventional DSLM, capable of illuminating the specimen holder. Finally, two cameras, one at the rear of the setup (in the optical path of the light-beam), used for pattern inspection and cuvette positioning, and one orthogonal to the specimen chamber for light sheet imaging, allowed direct observation from different angles. A filter wheel, equipped with a set of compatible filters, was placed in the path of the light sheet imaging camera to enable real fluorescence microscopy. All elements of the device are pictured in Figure 1b. Further details to the theoretical principles of light sheet bioprinting can be found in the supplementary information (Figure S11).

**Fig. 1.**
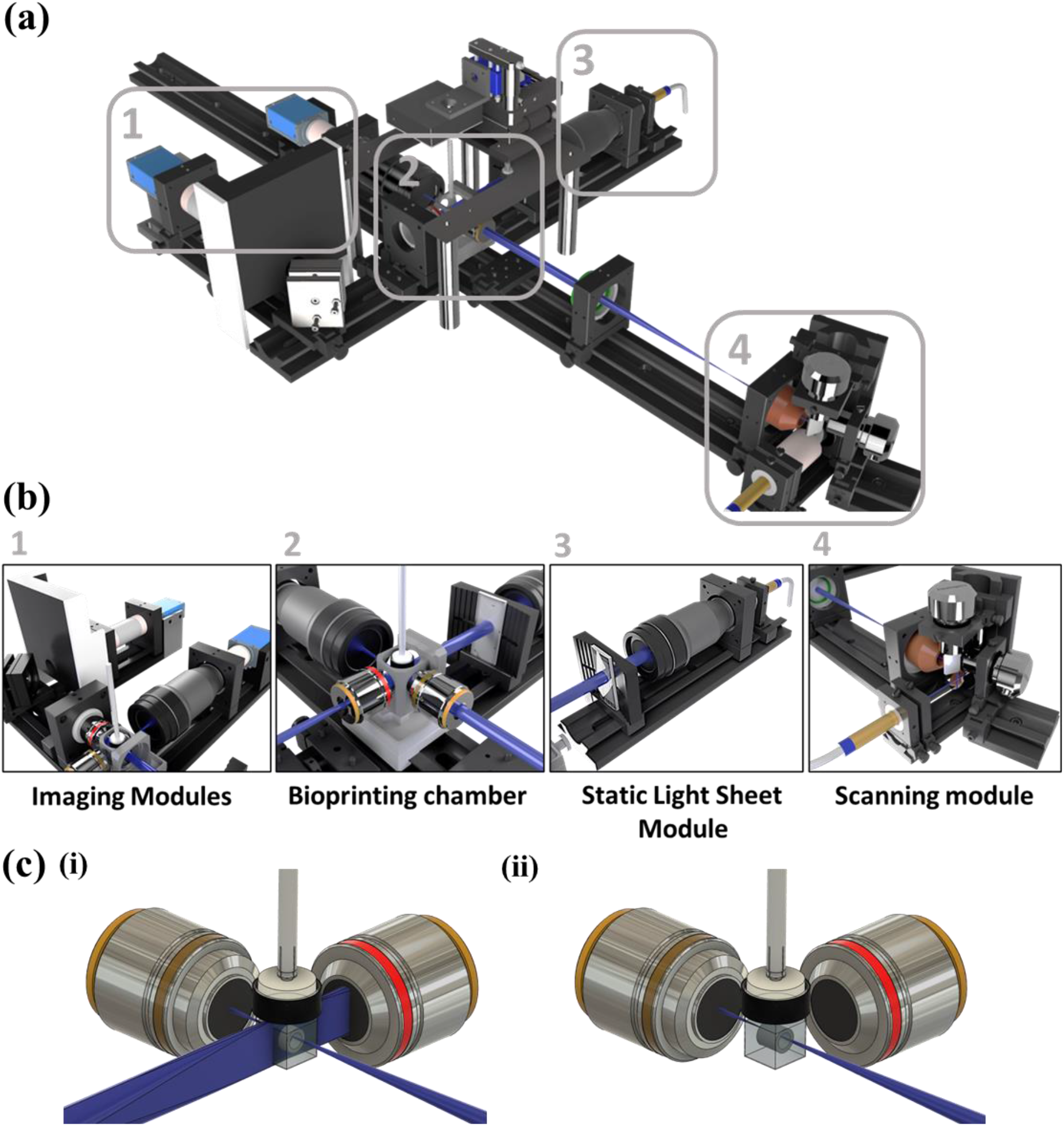
Overview of the light sheet bioprinter setup. (**a**) The optical set up for the light sheet bioprinter incorporates a patterning beam, a static orthogonal light sheet and imaging modules. (**b**) Overview of the light sheet patterning bioprinter. The bioprinter consists of four distinctive modules. (**b1**) The imaging module is capable of capturing patterns during the bioprinting process as well as fluorescence images before, during and after bioprinting. (**b2**) The bioprinting chamber is holding deionized water steady at 37°C to guarantee optimal conditions for cells and printing properties. (**b3**) The static light sheet is generated by a laser coupled with a beam expander and a cylindrical lens. (**b4**) The scanning module consists of three mirrors, one 45° mirror to introduce the laser beam into a galvanometer scanner pair, each one dedicated to scan the beam in a single axis (x and y). (**c**) At the focal point of the (scanned) laser beam and the static light sheet, a cuvette made of FEP-foil is holding bioink (hydrogel and cells) for the photocrosslinking process and imaging. (**i**) A double illumination or (**ii**) single laser beam crosslinking is possible for different printing requirements.

The bioprinting setup described in this study uses G-code, a widely used programming language for computer numerical control machines (*27*). G-code commands contain the type of action the device should perform (motions and positioning, turning on and off the laser, laser intensity) and the specific locations on the x-, y- and z-axes. The device then interprets these commands to move the galvanometer scanners and the z-axis of the stage with a defined speed and laser intensity to create a 2D pattern that, through mechanical motion of the cuvette in the z-axis, generates the 3D structure. The bioprinter reported here used a self-developed firmware written in C++ together with a controller software written in C#, steering every electronical device through a microcontroller. G-code files were uploaded through the controller software to the microcontroller and could subsequently be interpreted by the firmware. For this purpose, the G-code file was scanned line-by-line for type of action commands and the respective localization data. If a print command (‘G-command’) is found, the laser was turned on with a pre-defined power, and the galvanometer scanners moved the beam from a notional point A to point B, which were defined by xy-axes coordinates. After every line in a layer was scanned, a z-axis coordinate triggered the stage to move to the position of the next layer. This process was repeated for the whole length of the G-code file and automatically stopped the printing process once a stop command (‘M00’) was read.

Printing with only the laser beam could achieve high resolution results, provided the 3D object to be printed did not have complex internal structures located in the beam path. In this case, the power of the laser beam could be increased so that the light penetrates deeper into the cuvette and crosslinks several layers simultaneously (as in Figure 1c ii). This also led to a faster printing time. Supplementary Movie S1 shows the single beam patterning of a resolution wheel in real-time (similar to the one pictured in Figure 2a i). The increase in the laser power would however overexpose the first planes of the printed object.

**Fig. 2.**
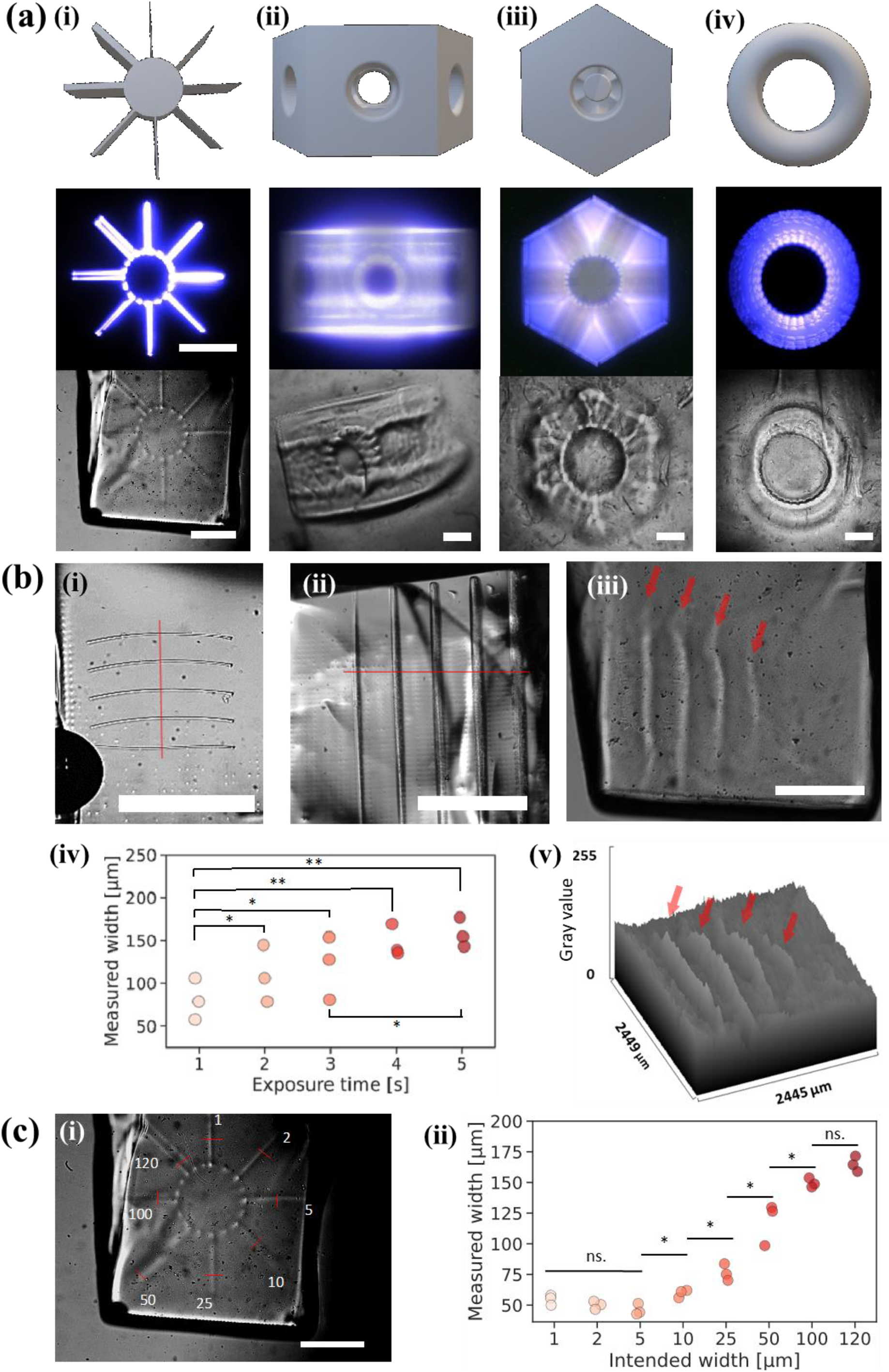
3D bioprinting of complex objects is accurate when using the light sheet bioprinting system. (**a**) Complex objects printed with the light sheet bioprinter show high resolution. Scale bar in light pattern and brightfield pictures (applies for all pictures): 1 mm. (**i**) Wheel of resolution (crosslinked in Cellendes hydrogel 1). Each branch of the wheel is a different thickness to show lateral resolution (xy-resolution): from top to top-left clockwise: 1, 2, 5, 10, 25, 50, 100 and 120 µm. The lowest successfully crosslinked thickness is 46.1 µm ± 4.6 µm. (**ii**) Side view of a liver lobule object. Side holes are 1.2 mm in diameter. (**iii**) Top view of the liver lobule. The edges are well defined. The diameter of the central hole is 2 mm. (**iv**) Print of a torus. The diameter and thickness are accurate. Additionally, the shape is overall smooth, which is difficult to achieve with extrusion bioprinters. The liver lobule and torus were printed with the GELMA/PEGDA hydrogel. (**b**) (**i**) The theoretical minimal axial crosslinking resolution of 11 µm for the laser beam is nearly achieved with 15.7 µm (mean) by using a resin for photocrosslinking and printing five lines with the lowest laser settings leading to photocrosslinking). (**ii**) A minimum printing resolution for the static light sheet was found with 80.8 µm when using a resin and (**iii**) around 80 to 158 µm when using the hydrogel. (**iv**) The static light sheet produces structures that are ranging from around 80 to 158 µm in thickness, depending on the laser power, using hydrogel. (**v**) A surface plot of the light intensity (8-bit) shows the crosslinked sheets protruding from the surface. Scale bars: 1 mm. (**c**) (**i**) The resolution wheel printed with hydrogel resolved structures of down to 42.8 µm, (**ii**) with spokes usually 1.3 – 2.6 times larger than intended, for spokes larger than minimum achieved resolution. Red arrows show the individual crosslinked sheets and red lines show the spots where the measurements for the width was conducted.

Therefore, for a more precise z-resolution, a static light sheet (405 nm) was introduced by a single convex lens to orthogonally illuminate the bioink-laden cuvette. By spatially defining the printing plane and using a second light source, the photocrosslinking threshold of the bioink was only exceeded where both illumination sources (static light sheet and laser beam) were superimposed. Hence, the bioprinted volume was limited to a voxel which size was determined by the width of the laser beam (xy) and the width of the static light sheet (z) (see Figure 1c i). Supplementary Figure S1 and Supplementary Figure S2 as well Supplementary Movie S2 show the static light sheet in the bioprinter set up.

Customizable cuvettes with dimensions from 1.5x1.5x2 mm³ to 10x10x12 mm³ (width x length x height) made of fluorinated ethylene propylene-foil (FEP-foil) based on previous work from Hötte et al. (*28*), were optically ideal vessels for bioprinting as well as for microscopy as the refractive index is close to the one of water (FEP n = 1.34; water n = 1.33, Supplementary Figure S3). Additionally, 3D bioprinted constructs could be kept in the same cuvette for later imaging and allowed for a streamlined process without unnecessary handling of the specimen. To create a cell-friendly environment, a custom 3D printed specimen chamber was designed, which incorporates a heating foil and a temperature sensor. The heating foil and the sensor were connected to a temperature regulator, which ensured an incubation of the cells at 37° C.

Various objects were printed with the bioprinter. To design, pattern and obtain an accurate three-dimensional object, a workflow was developed as demonstrated in Supplementary Figure S4. First, a 3D structure is modelled either by designing it in CAD software or by downloading an appropriate file from the internet (e.g., thingiverse.com). The exported ‘.stl’ file was sliced into lines and layers by a slicing software, resulting in a G-code file. A custom Python script was applied to the G-code to automate changes in such as speed and laser power. The adapted G-code file was then uploaded to the bioprinter software. To determine the accuracy of the 3D pattern, the rear camera recorded the individual illuminated planes of the structure, which could be assembled afterwards into an average or maximum intensity projection. After printing with the hydrogel, the rear camera was used to take high resolution photos or videos of the final construct. Constructs could then either be extracted from the cuvette for cultivation or kept in the same vessel and imaged with the light sheet microscopy function of the bioprinter, as will be described further.

### The light sheet bioprinter produces complex structures

Although light sheet properties have been well studied, the photocrosslinking properties of a light sheet are not yet determined. To understand the capabilities of the device, objects of different widths and depths were printed. In Figure 2a, the designed CAD model, the maximum intensity projection of the light pathway and the resulting object, are showcased as examples of the capabilities of the bioprinter. The laser-patterning took place in a thiol-ene photocrosslinkable hydrogel composed of a dextran-based backbone and a hyaluronic-acid crosslinker (Cellendes hydrogel 1, Table S5). First, the resolution wheel was demonstrated (Figure 2a i). The wheel was designed to have “spokes” of different thicknesses sprouting from the solid core (cylinder, designed to anchor the spokes and prevent them from collapsing). The illumination pattern showed that some spokes received more light than other, possibly leading to over-crosslinking and larger width than expected. The spokes were investigated further below. Next, a more complex object was printed, a liver lobule (Figure 2a ii and iii). As seen on the CAD image (first row) and the illumination pattern (second row), the object was meant to contain several hollow tunnels in all directions (x-, y- and z-axis). After printing, the structures are identifiable in the brightfield image of the object using transmission light (third row) which indicated that the resulting object was adequately crosslinked. Finally, a flat torus was designed and photocrosslinked (iv). A torus features various types of Gaussian curvatures which lead to different cell morphologies (*29*). The torus was accurately printed. The above-described objects were extracted from the cuvette after imaging and photographed in air under a stereomicroscope (see Supplementary Figure S5)

Next, the resolution was measured using a resin (Anycubic clear) that allowed printing of stiffer objects that could easily be imaged due to the higher refraction difference between not crosslinked and crosslinked resin (Figure 2b i, ii and iv). A single light sheet was printed by moving the light-beam once in the x-axis, and the length and width were measured to determine the axial (xy-) and lateral (z-) resolution, respectively as seen on Figure 2b i and Supplementary Figure S6. The theoretical minimal axial resolution of 11 µm (beam diameter at focal point) was almost met with 15.7 µm ± 9.1 µm (standard deviation) on average (median: 14.1 µm). Then, the resolution of the orthogonal light sheet was tested with the resin (Figure 2b ii and iv). The lowest setting on the laser engine was used (0.2 mW) while gradually decreasing the exposure time from five seconds to one second. It was noticed that the width of the photocrosslinked sheet decreased in a linear fashion with decreasing intensity. These results are in accordance with the Beer-Lambert law, where the intensity of the light decreases linearly in the z-depth (*30*) and seems to compensate for the absorption. Next, the Cellendes hydrogel (Table S7) was used. The hydrogel, as a softer extracellular matrix suitable for cell attachment and growth, is not as efficient in the crosslinking process as hard resin. The axial resolution was again tested using a resolution wheel (Figure 2c i). The spokes were ranging from 1 µm to 120 µm. All spokes are identifiable, which indicated successful photocrosslinking. Yet, the minimal observed resolution lies at 46.1 µm ± 4.6 µm on average for the 5 µm spoke (Figure 2c ii). The light pattern could potentially be at fault in the lack of accuracy: the current slicer was configurated to have a 5 µm light-beam (minimum thickness). This meant that, to crosslink a 10 µm spoke, the light beam was travelling back and forth closely to one another, which could lead to over-crosslinking. To understand the lateral resolution provided by the orthogonal static light sheet in the hydrogel, single sheets with descending power intensity (2, 1.6, 1.2, 0.8 mW) were crosslinked (Figure 2b iii and v and Supplementary Figure S7). The average width of the crosslinked light sheets for 0.8 mW was measured to be 178.2 µm ± 46.2 µm (standard deviation) and the thinnest crosslinked light sheet with 0.8 mW was measured to be 80.4 µm.

### Quality control of the bioink can be performed throughout the bioprinting process

The production of bioprinted objects using patterned light and subsequent imaging of the constructs was demonstrated in the previous section. The accuracy of the design could be determined in real-time with the help of a camera placed in the optical path of the laser. In addition to the rear camera, a side camera was installed to take advantage of the light sheet imaging capabilities of the system. Using this novel addition, the cells could be monitored throughout the bioprinting process. Here, we aim at understanding the impact of the bioprinting process on the cells and the fluorescent molecules. An angiogenesis model is used; fibroblasts stained with a mitochondrial dye (Hs27-MitoTracker) were co-cultured with HUVEC expressing GFP (GFP-HUVEC) as spheroids (cell aggregates (*31*)) for 48 hours (2:1 ratio). The spheroids were then collected and mixed with the polymer solution (Cellendes hydrogel 2, Table S7) before bioprinting. The object selected for this purpose was a hollow cylinder (2.5 mm height, 2.5 mm diameter with 1.5 mm diameter hole, see Figure 3a iii and Supplementary Figure S8) that was bioprinted in a 3x3x3.5 mm³ cuvette at an intensity of 12 mW, using a single light beam. Indeed, a hollow cylinder guaranteed proper medium diffusion for optimal cell growth. The cells were imaged as a z-stack using the light sheet microscope in the same position before and after bioprinting (Figure 3a i and Supplementary Movie S3).

**Fig. 3.**
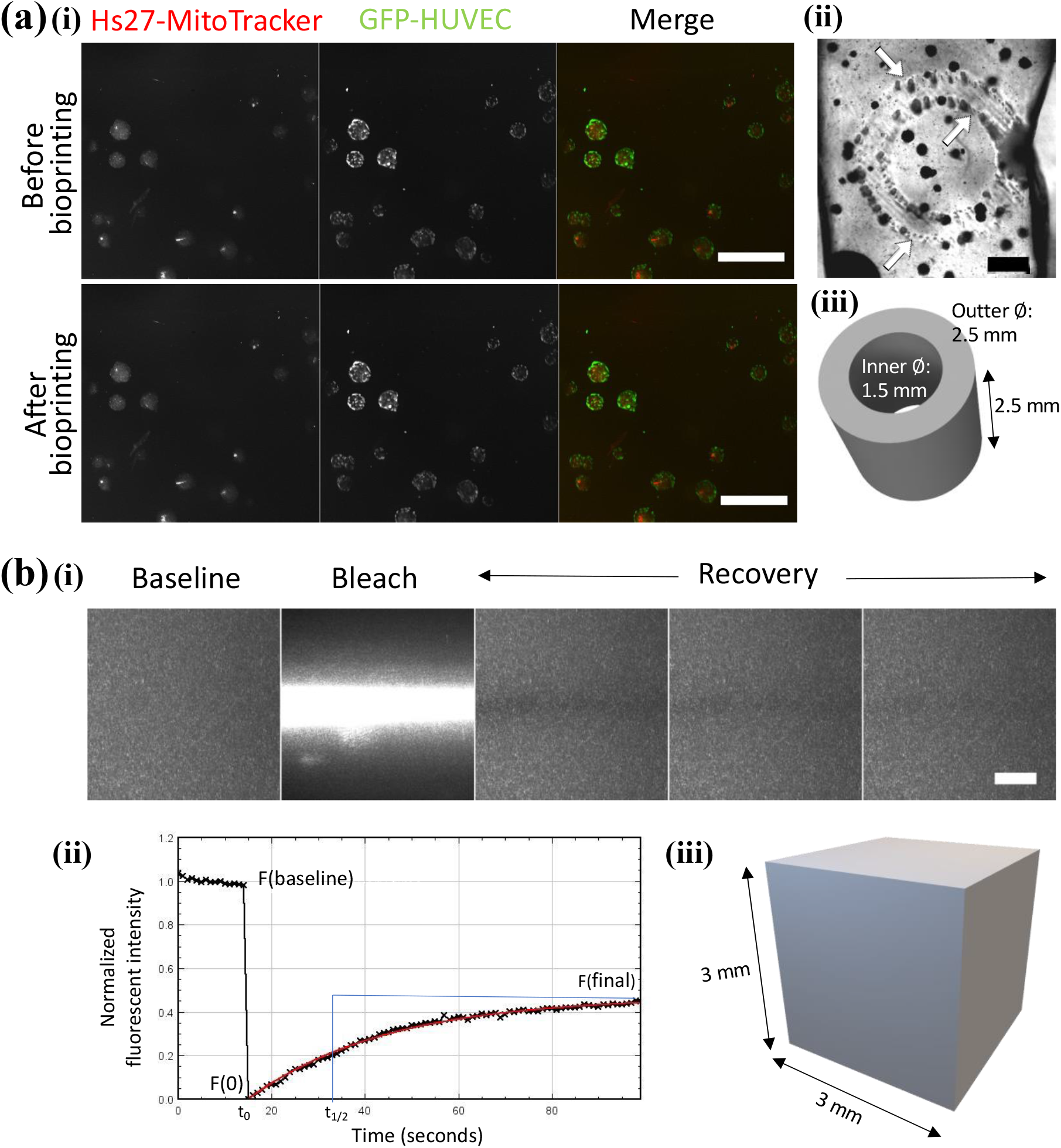
Streamlined imaging of the key elements in the bioprinting process (hydrogel and cells), for advanced quality control. (**a**) Bioprinted spheroids are imaged using a DLSM integrated in the bioprinting apparatus. Fibroblasts and HUVECs co-cultured as spheroids. (**i**) Hs27 cells stained with MitoTrackerRed and GFP-HUVEC cells were bioprinted. The intensity of the signal from Hs27-MitoTacker (in red) and GFP-HUVEC (in green) spheroids does not vary when imaged before or after bioprinting, the spheroids also did not change spatial positioning. Voxel size: 0.69 × 0.69 × 10 µm. Objective lenses: Zeiss A-Plan 2.5x/0.06 (excitation). Scale bar: 200 µm. (**ii**) The positioning of the cells or spheroids can be assessed by imaging the constructs in brightfield post-crosslinking. The boundaries of the printed objects are indicated by white arrows. Scale bar: 1000 µm. (**iii**) CAD rendering of the object selected for 3D bioprinting of cells (hollow cylinder). Printing intensity: 12 mW. (**b**) FRAP used in the bioprinter setting to assess crosslinking in the hydrogel. (**i**) Example of selected slices in a z-stack acquired during a FRAP experiment on a crosslinked hydrogel. First, a baseline is imaged (30% laser intensity, 10 images every second), then bleaching was performed (100% laser intensity, 10 seconds) before imaging the recovery diffusion though the bleached zone (18 mW, 100 images every second). The bleach zone is slowly repopulated with neighboring FITC-dextran molecules, eventually reaching a plateau. (**ii**) The fluorescent intensity of the bleached zone is normalized to a non-bleached zone and plotted against time. The half recovery time (t1/2) is calculated using the curve-fitting parameters. Note here that the photobleaching was not taken into consideration in the analysis. (**iii**) Rendering of the CAD file used for the FRAP experiment: 3x3x3 mm cube. Printing intensity: 18 mW.

It was noticed that the endothelial cells positioned themselves on the edge of the cell aggregate whereas the fibroblasts were compact in the core of the spheroids. No difference was noted between the before and after picture – the same parameters as for the light sheet (intensity, exposure time) were used, yet the fluorophores did not seem affected by the intensity of the beam during printing (no photobleaching). Additionally, the spheroids were situated locally identically which indicated that the hydrogel did not contract or expand during the bioprinting process. The rear camera was also used to check the placement of the light patterns on the hydrogel (not displayed) and the final product post-bioprinting (Figure 3a ii). The sharpness of the cylinder design is clearly identifiable from the front and the side, which indicated that the spheroids, albeit being tight spheres of highly mismatched refractive index, do not significantly affect the resolution of this object. Furthermore, the placement of the spheroids was observed in relation to the printed object. The spatial positioning of the cells to ensure the alignment with bioprinting designs is crucial, for stem cell niches or in the tumor microenvironment for instance. The number of encapsulated spheroids within the final product could be assessed to determine the efficiency of the bioprinting process. It is important to note that the use of light sheet microscopy allow for live imaging of non-cleared objects.

The bioprinted biological materials only represents one part of the process. The hydrogel plays a major role in the final bioprinted construct – the degree of crosslinking of the hydrogel, either determined by the concentration of the polymers or by the light intensity, affects the behavior of the cells and the diffusion of signals within the hydrogel (*32–34*). The crosslinking of the hydrogel depending on light intensity, exposure time, and other factors can be monitored using fluorescence recovery after photobleaching (FRAP). FRAP has been used as a method to determine the microstructure of hydrogels using the diffusion of fluorescent dyes (*34–36*). Confocal microscopy is commonly used to image FRAP results. However, light sheet microscopy, using orthogonal imaging, provides an additional view of the side diffusion that could prove be useful, for example if monitoring a hydrogel containing a stiffness gradient. As a proof of concept, a hydrogel (Cellendes hydrogel 1) containing a fluorescent dye (FITC-dextran 20 kDa) was crosslinked using the light-patterning system (a simple cube filling the volume of the cuvette was crosslinked, see Figure 3b iii), before analyzing the diffusion of the dye using FRAP.

Figure 3a i shows the three phases of FRAP. First, a baseline is recorded using a scanning light sheet, measuring the fluorescence level before bleaching. Next, a high intensity single beam was shone through the hydrogel to bleach the dye in a specific zone in the center of the image. Finally, the recovery, meaning the return of the fluorescent molecules to the bleached area, was imaged and the fluorescence was measured at a regular interval until a plateau was reached (Figure 3b i and ii). The time necessary to reach this plateau (which did not necessarily equate to the original baseline intensity) was calculated and half this time (half recovery time) was used as a conventional value to indicate the diffusion speed of the molecules. The lower the half recovery time, the faster the molecules would diffuse through the crosslinked hydrogel, indicating a looser network. Using this method, the user can therefore determine the necessary intensity to crosslink the hydrogel partially or fully.

All these quality control steps can be streamlined within the bioprinting process: the imaging and bioprinting actions are taking place in the same sample holder, in the same position, which removes the need for additional steps such the transfer of the object onto a well plate. This setting could furthermore be used in future projects to image the cells across a longer span of time (time lapse imaging).

### Light sheet bioprinting produces full-thickness skin tissues

Bioprinting is an inherently strenuous process for the cells. After passaging, the immersion in a synthetic polymer solution for an extended period while being processed through various methods are factors that influence the cell viability. For example, it has been shown that cells printed using syringe-based bioprinting lose viability due to the shear force produced by dispensation through a nozzle (*37–38*). Likewise, light-based bioprinting comes with hurdles for the cells. One factor that could influence cell viability is the wavelength used for photocrosslinking. Visible light is preferred to ultraviolet (UV) light which causes cell damage (*38–40*). Another aspect that could influence the cell viability is the presence of free radicals in the not crosslinked hydrogel. The chemical reaction of photocrosslinking involves cleaving a photoinitiator into two radical entities which trigger the chemical reaction (*41*) (for example, methacrylation or thiol-ene). The radicals, in contact with the cells, can create oxidation and cell damage (*42,43*).

Cell viability is therefore an indicator of the status of the cells after the bioprinting; measured by quantifying the number of dead cells over the total amount of cells. Human fibroblasts (Hs27 cells) were encapsulated in the Cellendes hydrogel 2 and bioprinted as a hollow cylinder, similarly to what was done in the previous section (laser intensity: 12-20 mW), to mimic a simplified dermis tissue. A live dead assay was performed directly after bioprinting (day 0) and after seven days in culture (in a well plate, immersed in medium). When measured directly after bioprinting, the average cell viability of the fibroblasts was high: 90% ± 8.98% (standard deviation or SD, Fig. S9). After seven days in culture, the viability remained important, with an average of 83% ± 4.34% (SD), proving that the bioprinting process and subsequent culture in a bioprinted hydrogel did not affect the cell viability (Figure 4b i). To produce a more complex tissue, Hs27 human fibroblasts and HaCaT human keratinocytes were co-cultured in a full-thickness construct. The Hs27 cells were encapsulated in the Cellendes hydrogel 2, bioprinted as a hollow cylinder and after three days, the HaCaT cells were seeded on the surface of the construct. After an additional seven days, the constructs were cultured in an air-liquid interface (ALI). The cell viability was measured 41 days post-bioprinting to be 74% ± 13.25% (SD). This slight decline could be mitigated by adding more complexity to the 3D bioprinted system, such as vascularization. Additionally, a high variability between the biological replicates was observed. Nevertheless, there was no significant difference found between the different culture lengths and day zero ((Welch t-test (n=3 to 5), p=0.35 and p=0.40 respectively).

**Fig. 4.**
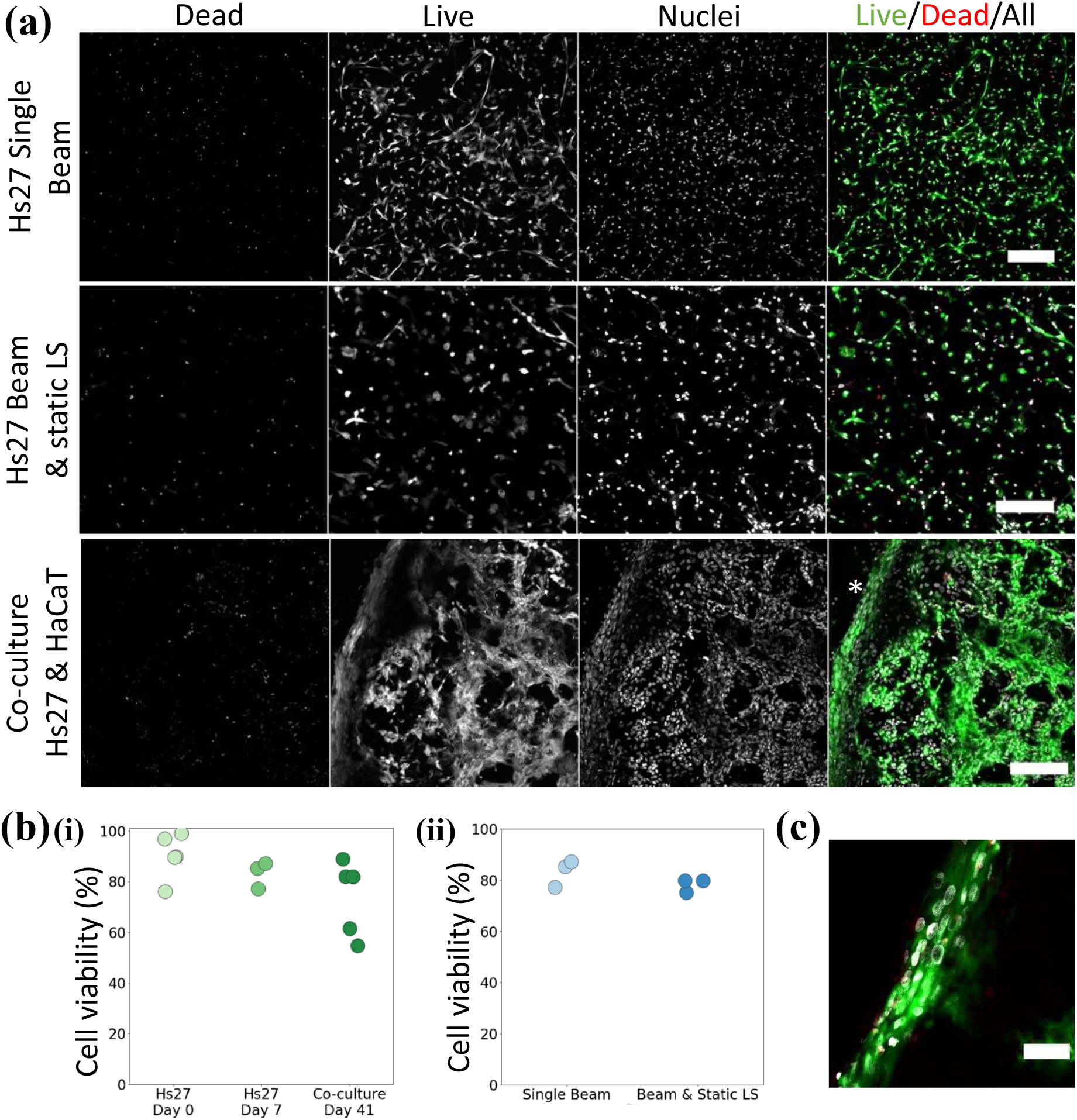
The viability of the cells bioprinted using the light sheet device. (**a**) Dead cells (stained with propidium iodide, PI), viable cells (stained with fluorescent diacetate, FDA) and overall cell population (stained with Hoechst 33342) of fibroblasts and/or keratinocytes encapsulated with a light sheet lithography device are imaged to determine the cell viability. The live cells are well spread out within the matrix. The image of the co-culture shows tight cellular structures present on the surface of the constructs that resemble an epidermis layering (indicated by an asterisk). (**b**) The images of FDA/PI/Hoechst-stained cells were segmented and analyzed to quantify the cell viability of different cultures of cells encapsulated with the light sheet lithography process. (**c**) Close-up of the outer layer of the bioprint that highlights the tight layer of cells, mostly alive. This image was extracted from slice 19 out of 57 of the z-stack and therefore shows a single layer of cells. Microscope: Zeiss AxioObserver LSM780. Objective: Plan ApoChromat 20×/0.8 M27. Voxel size “Hs27 Single Beam”: 0.52 × 0.52 × 6 µm. Voxel size “Hs27 Beam & static LS” and “Co-culture Hs27 & HaCaT”: 0.83 × 0.83 × 6 µm. Scale bar: 100 µm.

The impact of the addition of the static light sheet as was described previously was investigated. Fibroblasts Hs27 were printed in hollow cylinders (laser intensity 12 mW) with either a single beam or with the addition of the static light sheet. The viabilities of fibroblasts, cultured in immersion for seven days in constructs that were bioprinted with or without the use of the static light sheet were similar, with an average of 83% ± 4.34% and 78% ± 0.03% (SD), respectively (Figure 4b ii). Here again, no statistical difference was identified when comparing the single beam Hs27 culture with the static light sheet culture or with the long-term co-culture (Welch t-test (n=3), p=0.22).

When focusing on the edge of the long-term co-culture construct (marked on Figure 4a with an asterisk and as seen in the close-up on Figure 4c), a compact layer of cells (assumed to be keratinocytes) was visible. This layer seemed tight and somewhat stratified (although the uppermost cornified layer consisting of mostly dead cells is lacking). This structure resembled an immature epithelial layer as seen in vivo (*44*). To confirm the identity of the cell types and the physiological relevance of the bioprinted skin models, immunofluorescent staining of significant dermal and epidermal markers was completed.

The same objects, hollow cylinders, were bioprinted (laser intensity: 12 mW) and co-cultured in ALI conditions. First, the presence of vimentin was investigated. Vimentin is a cytoskeleton protein part of the intermediate filament family, which is highly expressed in fibroblasts (*45, 46*) which are predominantly found in the dermal part of the skin (*47*). Dermal fibroblasts are responsible for ECM production and hair follicle initiation (*48–50*). Vimentin was indeed present in most of the fibroblasts in a 3D bioprinted construct after seven days in culture and seemed to be expressed only in the elongated fibroblasts (Figure 5a). Likewise, collagen IV plays an important role as the main component of the basement membrane, the separation and support sheet-like structure between epidermis and dermis in the skin (*51, 52*). The fibroblasts, when cultured alone without keratinocytes, expressed collagen IV sporadically (Figure 5b). When seeding keratinocytes on top of the fibroblast-rich 3D construct and after culturing the bioprints for 41 days in ALI conditions (Figure 5c), the distribution of the proteins dramatically changed. Collagen IV was further expressed but was virtually covering the surface of the construct, which indicated formation of the basal membrane. The keratinocytes, identifiable by the expression of keratin 14 (*53*), were numerous above the basement membrane, although the tight layer of cells and beginning of stratification previously observed were not visible here. A possible explanation for the gaps in keratin 14 expression in the layer would be that some keratinocytes were keratin 14 negative, which might indicate keratinocyte differentiation (*53*). The distribution of collagen IV and keratin 14 can be further observed in cross sections of the object (Supplementary Figure S10).

**Fig. 5.**
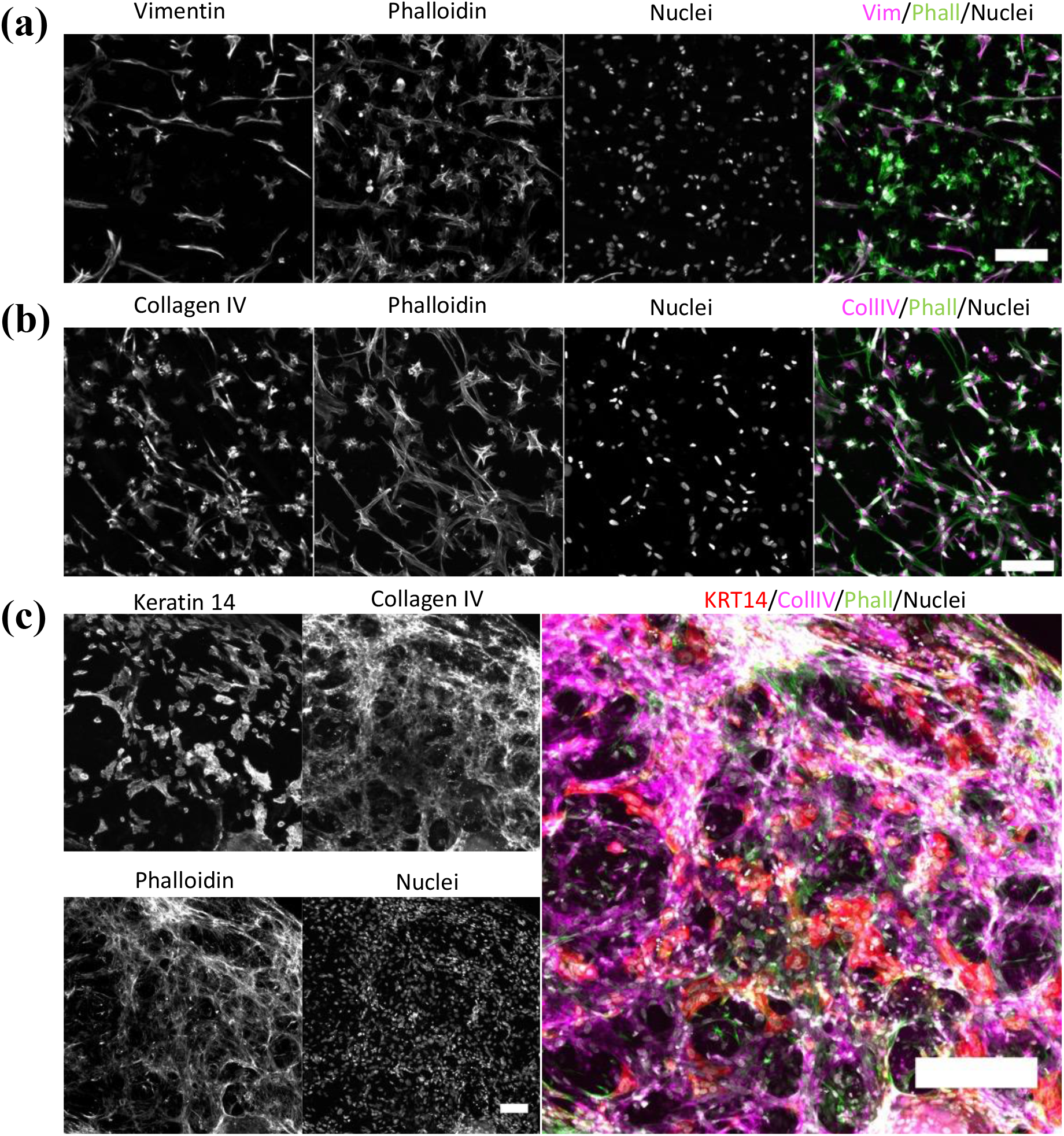
- Immunofluorescent staining of skin cells (Hs27 fibroblasts and HaCaT keratinocytes) cultured in 3D bioprinted objects display markers that are representative of dermis and epidermis. (**a**) Encapsulated Hs27 mimicking the dermis were stained against vimentin, phalloidin and Hoechst (respectively in pink, green and gray). Z-projection. Voxel size: 0.46 × 0.46 × 6 µm. Scale bar: 100 µm. (**b**) Further samples of encapsulated Hs27 were stained against collagen IV (CollIV), phalloidin (Phall) and Hoechst (respectively in magenta, green and gray). Z-projection. Voxel size: 0.42 × 0.42 × 6 µm. Scale bar: 100 µm. (**c**) A co-culture of Hs27 and HaCaT, representing a simplified epidermis-dermis model, were stained again Keratin 14 (KRT14), collagen IV (CollIV), phalloidin (Phall) and Hoechst (respectively in red, magenta, green and gray). Z-projection. Voxel size: 0.83 × 0.83 × 6 µm. Scale bar individual channels: 100 µm. Scale bar merge: 200 µm. Microscope: Zeiss AxioObserver LSM780. Objective: Plan ApoChromat 20×/0.8 M27.

The 3D bioprinted objects produced using the light sheet lithography device presented in the work were able to survive in a medium- (7 days) and long-term (41 days) culture and retained their main characteristics, namely elongation of the fibroblasts and invasion of the matrix, formation of a basement membrane and initiation of an epithelial layer.

## Discussion

Producing faster and high-resolution 3D bioprinting methods is a primary goal in the field of bioengineering since its inception. However, one should not come at the expense of the other, speed in particular should not come at the cost of resolution or design geometry. Additionally, the race for ever faster and high-resolution devices does not always account for an application-based point of view. Indeed, is a nanometer or micrometer resolution always necessary? The field of tissue engineering would certainly benefit from adding streamlines and user-friendly monitoring methods that go beyond the actual bioprinting process. We introduced in this work a novel bioprinting device that, using the principle behind light sheet microscopy, can produce complex structures while also combining an imaging device that can account for the cells and the hydrogel’s state at any time throughout the bioprinting process. Cells encapsulated in a photocrosslinkable hydrogel showed high viability post-bioprinting, even after long-term culture, and encouraging tissue-specific markers. The device described is versatile, in the way that it can be combined with other types of illuminating systems, such as two-photon, volumetric bioprinting or xolography 3D printing. While similar, these methods do not allow for as much flexibility. An application of xolography in the bioprinting field might be of interest due to the fast and high-resolution generation of 3D matter, however it was not shown yet and is possibly not straightforward due to the highly chemical nature (dual-color photoinitiator) and high radiation dosage in the UV-spectrum (375 nm) of the method. Light sheet bioprinting has been shown to use 20-50 times less optical energy (mJ/cm^2^) than volumetric printing, making it an attractive option for reducing optical energy usage in bioprinting applications, as excessive exposure to optical energy can potentially damage cells and tissues.

Improvements are nevertheless necessary to make this bioprinter ready for market. First, even though all components (medium, hydrogel polymers and sample holder) have been carefully selected to avoid refractive index mismatch, additional light scattering could be further minimized to improve resolution and design accuracy. For example, the use of a contrast agent to homogenize refractive index between cells and hydrogel (*55*) or the use of correction masks (*56*) are methods that could be of interest in conjunction with the light sheet lithography bioprinter. The fact that only one sample at a time can be currently bioprinted is an obvious roadblock to high-throughput drug discovery models; however, using an inversed light sheet setup, the technique could be adapted to accommodate well-plates.

So far, the method of “slicing” the CAD model of the object to be bioprinted is still the state-of-the-art in either extrusion or stereolithography methods. The slicing generates a stack of 2D images that are projected on the xy-axis or, in the case of volumetric bioprinting, along the rotation axis (radon transform of the CAD file). The resulting pattern is in these cases invariably a 2D projection. In our work, we showed that the use of a light beam of variable intensity, combined or not with an orthogonal static light sheet allows for more flexibility in terms of photocrosslinking geometries. The crosslinking across a plane is of course permitted, as we showed in this work, however single beam crosslinking or even point crosslinking, similar to what is done in two-photon polymerization, would be more desirable for faster and more accurate bioprinting of complex structures. To remedy this gap in the current technology, the next step would be to develop a slicer software capable of analyzing the structure to be printed and deducting the best method for photocrosslinking (plane by plane, single beam, single point, or a combination of those) and the laser intensity necessary for this application.

The ability to create gradients of stiffness across multiple planes, including the xy-axis in addition to the z-axis, would be a valuable tool for tissue engineering applications. This flexibility could enable the creation of more complex tissue structures with precise mechanical properties. Moreover, the versatility of the bioprinting device demonstrated in this study suggests that it could have broader applications beyond tissue engineering, such as hydrogel testing and drug discovery. Moving forward, the future of bioprinting lies in the development of more versatile machines that combine bioprinting, imaging, and quality control capabilities. The 3D bioprinter presented in this study shows great potential for such future developments, which could allow for the production of more functionally accurate tissues and organs.

## Materials and Methods

### Materials

The clear resin used for the resolution tests (3D printing UV sensitive resin, Basic, 1 kg) was purchased from Anycubic Technology Co. Limited. The porcine skin type A and methacrylic anhydride was purchased from Sigma Aldrich Chemie GmbH. The dialyzing membranes were obtained from Spectrumlabs. The freeze drier was an Alpha1-4LD from Christ and the spectrometer a DMX-500 high resolution NMR spectrometer from Brucker. The polyethylene glycol diacrylate (PEGDA), Lithium-Phenyl-2,4,6- trimethylbenzoylphosphinat (LAP), tartrazine and FITC-dextran were purchased from Sigma Aldrich Chemie GmbH. The phosphate buffer saline (PBS) was purchased from Gibco, ThermoFisher Scientific. All hydrogel components for the Cellendes hydrogel were provided by Cellendes GmbH as part of the BRIGHTER project. The Hs-27 human foreskin fibroblasts were purchased from the American Type Culture Collection (ATCC, CRL-1634). The HaCaT human keratinocytes were purchased from Cell lines services (CLS, 300493). Green fluorescent protein-expressing human umbilical vein endothelial cells (GFP-HUVEC) were purchased from Pelo Biotech (cAP-0001GFP). Media, supplements and cell culture consumables were purchased from ThermoFisher Scientific. Medium and supplements for the endothelial cells as well as the flask speed coating solution were purchased from Pelo Biotech. Normocin was purchased from Invivogen.

The cell culture plate inserts (transwells) for 24 wells (PET membrane, 3.0 µm pore size) were purchased from VWR International.

Fluorescein isothiocyanate–dextran (FITC-dextran) was purchased from Sigma Aldrich Chemie GmbH. The antibodies and dyes were purchased and diluted according to Table S4. Paraformaldehyde (PFA) and triton X-100 were purchased from MilliporeSigma, glycine, tween-20 and albumin fraction V (BSA) were purchased from Carl Roth GmbH. Goat serum was purchased from ThermoFisher Scientific.

The overall pictures of the bioprinted objects were taken using the Zeiss SteREO Discovery V8 stereomicroscope (Carl Zeiss GmbH). The cell viability and immunofluorescent staining pictures were taken using the Zeiss AxioObserver LSM780 confocal microscope (Carl Zeiss GmbH).

### Bioprinter setup

Optical parts were installed onto an optical breadboard, using the OWIS 45 and 65 rail system.

The multi-wavelength iChrome CLE-CD laser engine used was purchased from TOPTICA Photonics AG. It comprises four wavelengths (395/60; 488/20; 561/20; 640/20 nm/mW) in one engine. Another iChrome CLE laser engine with four wavelengths (405/20; 488/20; 561/20; 640/20 nm/mW) was used together with a zoom beam expander (1x − 8x, S6ASS2075-067, Sill Optics GmbH & Co. KG) and a cylindrical lens (f=120 mm) for creating the static light sheet. Two DynAxis 3S galvanometer scanners (one for x- and one for y-axis) were purchased (SCANLAB GmbH) together with their respective controller boards. A telecentric f-theta lens (f= 40 mm), specifically manufactured for the use with near-UV light, was purchased from Sill Optics GmbH & Co. KG.

Objective lenses with 2.5x (EC Epiplan-Neofluar 2.5x/0.06, M27, WD: 15.1mm) and 5x magnification (Plan-Neofluar 5x/0.16, M27, WD: 18.5 mm) from Carl Zeiss were used for illumination and detection, but are easily replaceable by other objectives with e.g., higher or lower magnification and numerical aperture. A tube lens (Carl Zeiss, 1x, f= 164.5 mm) was used to create a real intermediate image before the light enters the objective lens. A PIFOC objective scanner (Physik Instrumente, P-725.4CD) together with a compatible controller (Physik Instrumente, E-709) was used for focusing the illumination objective.

Three M-111.2DG1 compact linear stages (Physik Instrumente) were coupled with a M- 116 360-degree precision rotation stage (Physik Instrumente) to allow a movement of the cuvette in four axes. A C-884 DC motor (Physik Instrumente) controller was used for steering the stages. Two 4k resolution cameras from The Imaging Source Europe GmbH were purchased for pattern observation and cuvette positioning (DFK33UX34) and for light sheet image detection (DMK33UX34). Another zoom beam expander (1x − 8x, S6ASS2075-067, Sill Optics GmbH & Co. KG.) was used to focus light into the pattern observation camera and another tube lens (Carl Zeiss, 1x, f=164.5 mm) was placed in front of the light sheet image detection camera. A computer-controlled filter wheel and its corresponding controller (Sutter Instruments, Lambda 10-3) equipped with four filters were used to filter out non-fluorescent signals for the light sheet imaging. Light is directed into the light sheet imaging camera via a round protected silver mirror (Thorlabs, Ø1").

The specimen chamber was custom designed, and 3D printed on an Anycubic Photon Mono X using black resin (Anycubic). The chamber includes windows made of either cover glass (illumination) or FEP-foil (detection) and an insert for a temperature sensor and a heating foil, which can be controlled via a temperature regulator (Winkler, WRT- 2000). Stainless steel stage holders and specimen holders were machined in-house and equipped with a magnetic head for seamless attachment to the stage.

A custom-built PCB based on an Arduino clone (PJRC, Teensy 4.1), was used to centrally connect, and control the laser units, galvanometer scanners, stages, cameras, and filter wheel. Custom digital-to-analog converter boards were used to address analog inputs on some devices (laser units, galvanometer scanners). Custom digital-to-serial converter boards were used to address serial inputs on other devices (stage controller, PIFOC controller).

### Bioprinter handling and software

A custom firmware, flashed onto a Teensy 4.1 microcontroller and written in C++, was used for controlling the bioprinter and microscope components. Functions in the software were separated for the use of microscopy and bioprinting features. The main function for bioprinting is the interpretation of G-code files. The file was read line by line by the software and based on the type of action in the G-code (’M’ and ’G’ values) the software recognizes which hardware was addressed. Based on the localization data (xyz-coordinates) the software could perform the movement pattern of the hardware (galvanometer scanners, stage) and modulate the respective intensity and velocity settings based on the ’S’ and ’F’ values. Automatic camera exposure for one layer was set by using the ‘M219’ value and dwell time between image exposure by using the ‘P’ value together with a numerical value translating into milliseconds.

3D models were designed in the computer-aided design software Fusion 360 (Autodesk).

G-code files were generated by using slicer software, in this case Slic3r (https://slic3r.org/, version 1.3.0), an open-source programme was used. A self-written Python script was developed to allow for automizing the customization (‘S’ and ‘F’ values) of G-code files, which cannot be done in the slicer software. An additional feature of the script is the calculation of the total pattern track length, resulting in the total print time when divided by the scanning speed.

The sample holders used were adapted from Hötte et al. 2019 (*28*). The vacuum-formed ultra-thin fluorocarbon (FEP) foils cuvettes were adapted into 3 or 10 mm (length and width), so larger objects could be bioprinted. The molds for thermoforming were designed on Fusion 360 (Autodesk) and printed on 3D printers of the Anycubic Photon series (Anycubic).

Laser power for the single beam (Table S1) and the static light sheet (Table S2) were measured at the focal points and subsequent calculations for each 3D (bio-)printed construct are listed in Table S3.

### Preparation of photocrosslinkable hydrogels

The GelMA/PEGDA hydrogel was composed of 10% w/v gelatin methacrylate (GelMA around 80% bloom) and 10% w/v polyethylene glycol diacrylate (PEGDA average Mn 4000) mixed with 0.2% w/v LAP and 0.025% w/v tartrazine (Table S6). The gelatin methacrylate was prepared following a protocol adapted from Loessner et al. 2016 (57–59). Briefly, gelatin from porcine skin type A was dissolved in PBS at 50°C under stirring conditions for 2 h to obtain a 10% (w/v) gelatin solution. Methacrylic anhydride (MA, 5% v/v) was added at a rate of 0.5 ml min^-1^ and the mixture was left under stirring conditions for one hour. Then, after centrifugating the solution (1200 rcf for 3 min), the reaction was stopped by adding Milli-Q water to the supernatant. The resulting mixture was dialyzed using 6–8 kDa of molecular weight cut-off (MWCO) membranes (Spectra/por) against Milli-Q water at 40°C, replaced every four hours for three days. The pH of the dialyzed products was subsequently adjusted to 7.4. The samples were kept overnight at −80°C and lyophilized for 4 and 5 days using a freeze drier. The degree of methacrylation was inspected using nuclear magnetic resonance (NMR) spectrometry (60). GelMA and PEGDA with LAP were separately mixed with PBS at 65°C for two hours then were combined, tartrazine was added and the mix was left at 37°C for an additional hour.

The Cellendes hydrogel was composed of two precursors: a main polymer (dextran (Dex)) carrying norbornene thiol-reactive group (N-Dex), and a thiol-containing crosslinker (with a backbone of polyethylene glycol (PEG-Link). The precursors were additionally functionalized to provide a cell-friendly environment when encapsulating cells. A cell-adhesion motif (arginyl-glycyl-aspartic acid or RGD) had been added by the supplier to the main precursor (RGD-N-Dex) while a cell-degradable, matrix metalloproteinase sensitive peptide (CD) had been added by the supplier to the hyaluronic acid crosslinker (CD-HyLink). The final concentration of norbornene and thiol was adjusted to achieve different degrees of crosslinking and thus various hydrogel stiffnesses. The details of the concentrations are listed in Table S5 and Table S7. The main polymer and the crosslinker were mixed with a HEPES-phosphate buffer without phenol red (pH 7,2), water and LAP before adding the cell suspension (where applicable). In addition, the pre-gel solution contained 0.1% low melting point (LMP) agarose. For gelation of the LMP agarose, the pre-gel solution was kept on ice for at least five minutes prior to bioprinting.

### Fluorescence recovery after photobleaching (FRAP)

The RGD-N-Dex and CD-HyLink bioink was used (Cellendes hydrogel 2). The water component was replaced by FITC-dextran diluted in water (20 kDa, 1 mg/ml). The hydrogel was placed in the sample holder and the bioprinting device was then used to crosslink a cuboid (3x3x3 mm³). The microscope part of the device (as previously described) was subsequently used to image the molecular diffusion of the FITC-dextran with a 488 nm beam. First, a baseline was imaged with a light sheet (10 images taken every second at 18 mW). Then, the light sheet height was lowered to zero and the intensity increased to 100% (60 mW) so that a single beam could be used to bleach an area of the field of view (10 seconds). Lastly, the post-bleach recovery was imaged using the light sheet scanning for 100 repetitions at 18 mW, every second.

The images were analyzed using Fiji by ImageJ (version 1.53c, U. S. National Institutes of Health). A Jython script developed by Johannes Schindelin (61) was used to extract the mobile fraction and half recovery time (t1/2), measured as follows:

Mobile fraction = (F(final)-F(0))/(F(baseline)-F(0))

t1/2 = F(final) – F(0)

With F(final) the final recovery intensity, F(0) the intensity at t=0 right after bleaching and F(baseline) the baseline intensity.

### Cell culture and encapsulation in the photocrosslinkable hydrogel

The cells were handled in sterile conditions and cultured in an incubator at 37°C and 5% CO2. The Hs27 cells and HaCaT cells in DMEM supplemented with 4.5 g/L glucose and 2 mM glutamine. Both media were also supplemented with 10% fetal bovine serum (FBS) and 1% penicillin/streptomycin (PenStrep). The GFP-HUVEC cells were cultured with the provided medium, supplements and antibiotics from Pelo Biotech. The cells were cultured in 25 or 75 cm² flasks, coated with the speed coating solution (Pelo Biotech), the medium was changed every two to three days and the cells passaged every week.

The hydrogel used to encapsulate cells was Cellendes hydrogel 2 (Table S7). To encapsulate the cells in the hydrogel before 3D bioprinting, the cells were detached from the flask using Accutase and collected by centrifugation in a pellet (300 rcf, 5 minutes). The supernatant was discarded and the cells were resuspended in the previously prepared hydrogel (see previous sections) with a density of 2 million cells/ml. The agarose was added (to keep the cells in suspension) and the cell/hydrogel mixture was kept on ice for at least 5 minutes or until photocrosslinking. The cell/hydrogel mixture was pipetted into the cuvette (the 3 mm cuvette contained 30 µl, the 10 mm cuvette contained 1000 µl) which was sealed and brought to the bioprinter. After bioprinting, the 3 mm cuvette was opened using a scalpel and the construct was extracted using a metal spatula (the 10 mm cuvette had a big enough opening to extract objects without cutting it open). The bioprinted objects were washed in PBS supplemented with 1:500 Normocin to prevent potential contamination linked to handling and are subsequently cultured in a well plate.

In the case of a Hs27 and HaCaT co-culture, the fibroblasts-rich construct was 3D- bioprinted as described above, introduced to the upper compartment of a transwell and subsequently incubated in the medium for 3 days. The HaCaT human keratinocytes were then passaged and the medium/cell mixture (1 million cells/ml, 400 000 cells/well) was pipetted on top of the bioprinted constructs. The immersed culture was maintained for an additional 7 days. Thereafter, the medium contained on the upper part of the transwell was removed while the medium in the lower part of the transwell remained, as is required in an air-liquid (ALI) culture. These conditions were maintained for 41 days with medium changes of the lower compartment every other day.

The Hs27 and GFP-HUVEC co-culture was performed by co-culturing the cells as spheroids in a Sphericalplate 5D well-plate (Kugelmeier Ltd). Each well contained 750 microwells. The spheroids were composed of 1500 cells and were a combination of 2:1 Hs27 to GFP-HUVEC. The Hs27 cells were incubated in MitoTracker Red CMXRos (ThermoFisher) for 15 minutes in a serum-free medium prior to the spheroid formation, as indicated in Table S4. The culture medium used for the co-culture was a mix of 50% Hs27 medium and 50% GFP-HUVEC. After 48 hours of culture in the Spherical plate, the spheroids were collected and encapsulated in the Cellendes hydrogel for imaging and bioprinting.

The specifications for bioprinting are included in Table S3. The energy dose required to bioprint the object (a hollow cylinder in the case of cell encapsulation) might vary on the volume of medium left with the centrifugated pellet. Although one tried to minimize the volume as much as possible, when the volume was high, the hydrogel was slightly diluted and the energy required for crosslinking needed to be higher. The energy ranged from 5.02 to 10.30 mJ/cm².

### Cell viability and immunofluorescence staining

The viability of cells after bioprinting was assessed using a propidium iodide (PI) and fluorescein diacetate (FDA) staining. The bioprinted constructs were extracted from the cuvette, washed with warmed PBS, then incubated at 37°C for 15 minutes in medium without supplements and phenol red, that contained 1:100 PI, 1:500 FDA and 1:500 Hoechst (nucleus stain). After incubation, the constructs were once more washed in PBS and imaged in medium.

The immunofluorescence staining followed a previous protocol (58). All the steps were performed at room temperature except otherwise indicated. Briefly, the bioprinted constructs were fixed in 4% PFA in PBS for 30 minutes, then washed thrice in PBS. Permeabilization followed using Triton X-100 (0.3% v/v) in PBS for 40 min before washing thrice in 0.1 M glycine in PBS and thrice in 0.1% Triton X-100 in PBS (PBS-T). The samples were subsequently blocked for 1 hour in a freshly prepared blocking solution (10% goat serum in BSA (0.1%), Triton X-100 (0.2%), Tween-20 (0.05%) in PBS). The primary antibodies (Table S4) were diluted in blocking solution and incubated at 37°C overnight. On the next day, the samples were washed in PBST-T thrice before incubating in the secondary antibody solution (also diluted in blocking solution) for 2 hours at 37°C. A final wash with PBS-T (three times) was performed before imaging in 2% penicillin/streptomycin in PBS. The list of antibodies and dyes is provided in the supplementary material (Table S4).

### Image processing and statistical analysis

Image processing was conducted in Fiji by ImageJ (62) (version 1.53c, U. S. National Institutes of Health). The images were cropped and brightness and contrast were adjusted. The images captured within the bioprinter were additionally deconvoluted using the PSF generator (63) and DeconvolutionLab2 (64) plugins. The data produced by the live dead assays and the immunofluorescent staining was processed using ImageJ. The images presented in this work are z-projections of the z-stacks imaged, unless otherwise specified. To quantify the live-dead assay data, the cells stained in the dead channel (PI staining) and the nuclei channel (Hoechst 33342) were separately counted. A gaussian blur filter was applied to images (radius 2.0), then an intensity threshold was applied so that a binary image of the cells was created. When necessary, a watershed algorithm was additionally used to separate adjacent cells. Finally, the 3D object counter plugin (65) was applied to count the number of cells segmented.

The statistical analysis was conducted on Python 3.9 (Python software foundation). The samples’ normality was tested with a Shapiro-Wilk test (p>0.01). Subsequently, statistical comparison between two groups was tested with Welch t-test (p<0.01). Exact p-value resulting from the tests are included in the text. Plots were generated on Python using the Pandas (66) Seaborn (67) and Matplot (68) libraries. Graphical abstract created using Biorender.com.

## Supporting information

Supplementary materials - Hafa et al. 2023

Supplementary Materials Hafa et al. - Movie S1

Supplementary Materials Hafa et al. - Movie S2

Supplementary Materials Hafa et al. - Movie S3

## Acknowledgments

The authors would like to thank all the collaborators of the BRIGHTER consortium: Gustaf Mårtensson, Helmut Wurst, Brigitte Angres and Ruby Shalom-Feuerstein. The authors thank Angela Cirulli for her assistance in testing the hydrogel preparation, and for the hydrogel crosslinking and the cell culture protocols. The authors also thank Sven Plath and the Mechanical Workshop of the Biological Faculty at Goethe Universität Frankfurt for helping with the assembly of the first prototype and for their continuous support throughout this project.

## Author contributions

FP and LH designed the bioprinter. LH assembled and tested the optical components, designed the patterning function and tested the software and electronics together with LRP. Controller software was written by LRP. CADs were designed by LB and LH and imaged by LH. LB and LH designed the experiments relating to resolution and LH conducted them. Hydrogel testing experiments (FRAP) were designed and conducted by LB. NT produced the gelatin methacrylate. EM and NT provided the cells, designed cell encapsulating protocols and conducted preliminary tests on the hydrogels and their compatibility with cells. LB conducted the experiments involving culturing, bioprinting, imaging, viability and immunostaining of cells. FP, LB and LH wrote the manuscript. Support to research was provided by EHKS. The research was conceived and supervised by FP. All the authors read and revised the manuscript.

## Data and materials availability

All data needed to evaluate the conclusions in the paper are present in the paper and/or the supplementary materials.

