## Supplementary materials - Hafa et al. 2023 for "Laser patterning bioprinting using a light sheet-based system equipped with light sheet imaging produces long-term viable skin constructs"

Levin Hafa *et al.*

**This PDF file includes:**

Supplementary Text
Figs. S1 to S11
Tables S1 to S8
Legends for Movies S1 to S3

**Other Supplementary Materials for this manuscript include the following:**

Movies S1 to S3

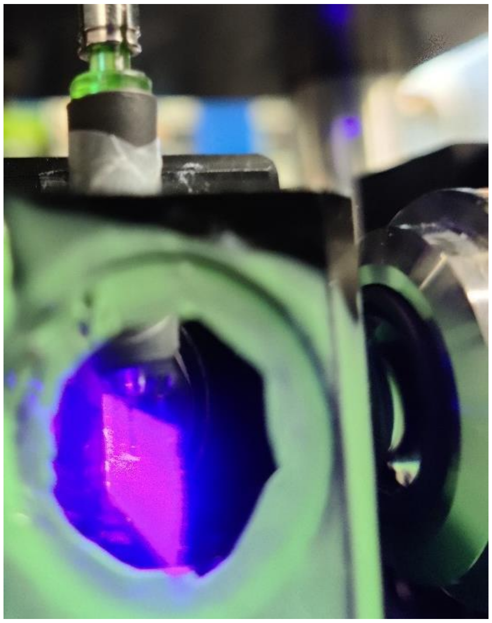

**Fig. S1. Picture of the static light sheet (view from the side), crossing the specimen chamber.**

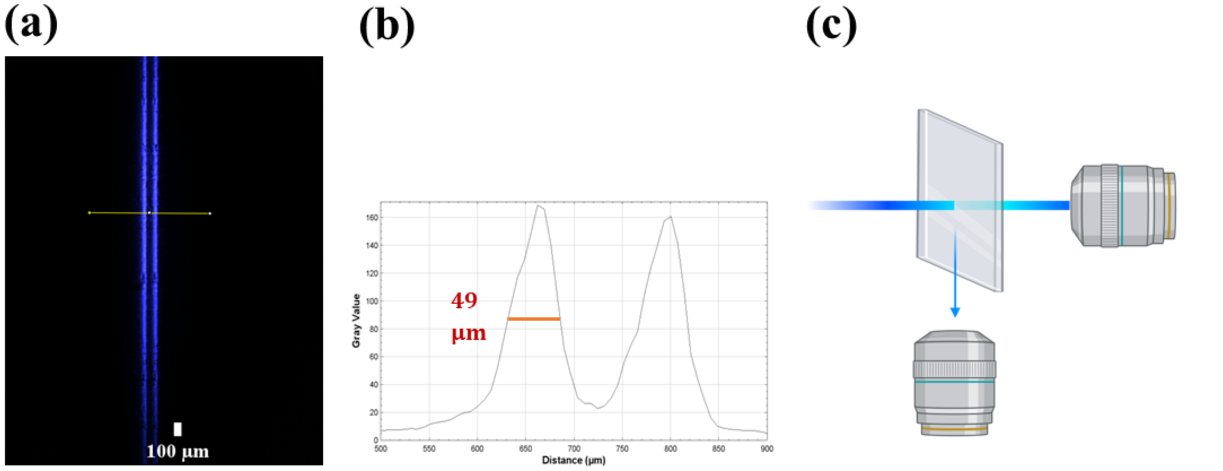

**Fig. S2. Measurements of the static light sheet.** (**a**) Light sheet microscope image showing the width for the static light sheet at 405 nm wavelength. (**b**) Full width half maximum (FWHM) measurement to define exact width of the light sheet. Measurement in the middle of the light sheet was found to be 49 µm. (**c**) The impression of a double light sheet is due to refraction from a cover glass.

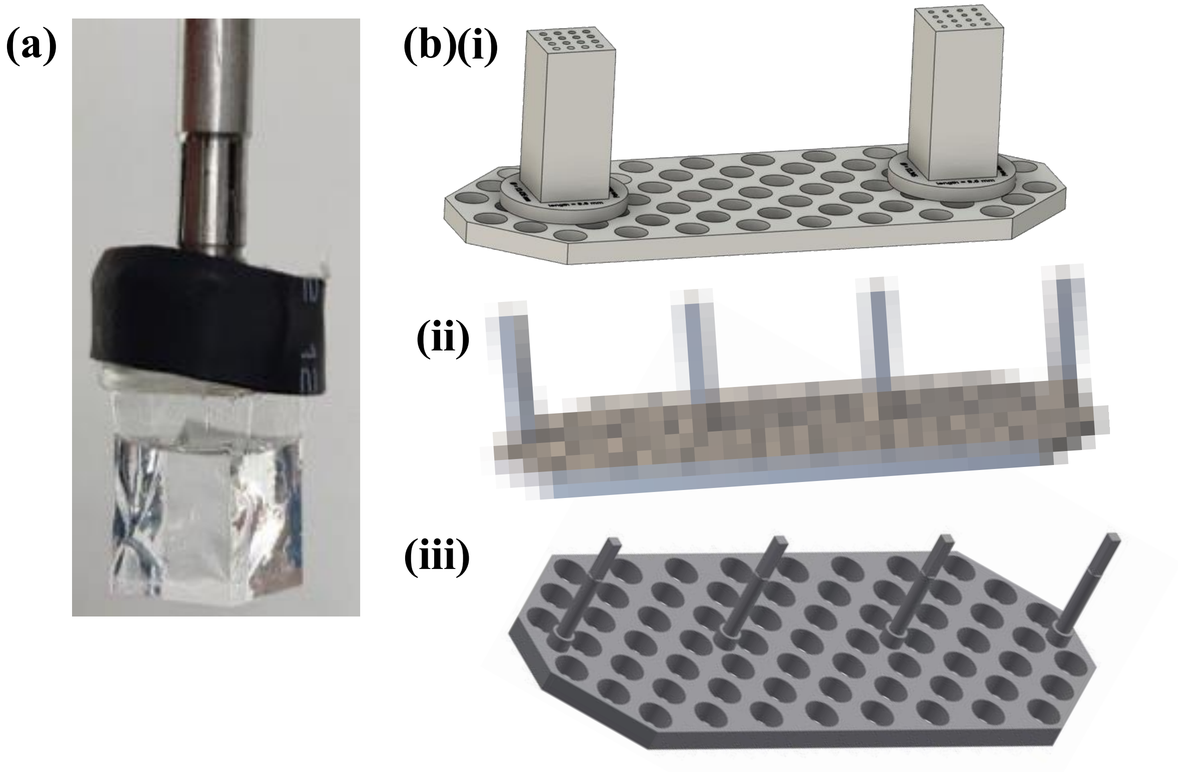

**Fig. S3. The cuvette used as a sample holder for 3D bioprinting with the light sheet device.** (**a**) Picture of the 10x10x10 mm model of the cuvette, filled with water. The FEP foil cuvette is connected to the sample holder metal rod through the addition of a shrinking tube (in black). (**b**) The cuvettes are produced using the thermo-forming method described by Hötte et al. (*28*). (**i**) Positive mold for the 10x10x10 mm cuvette. The holes are used to maximize the efficiency of the vacuum during thermoforming. (**ii**) 3x3x5 mm cuvette mold. (**iii**) 1,5x1,5x5 mm cuvette mold.

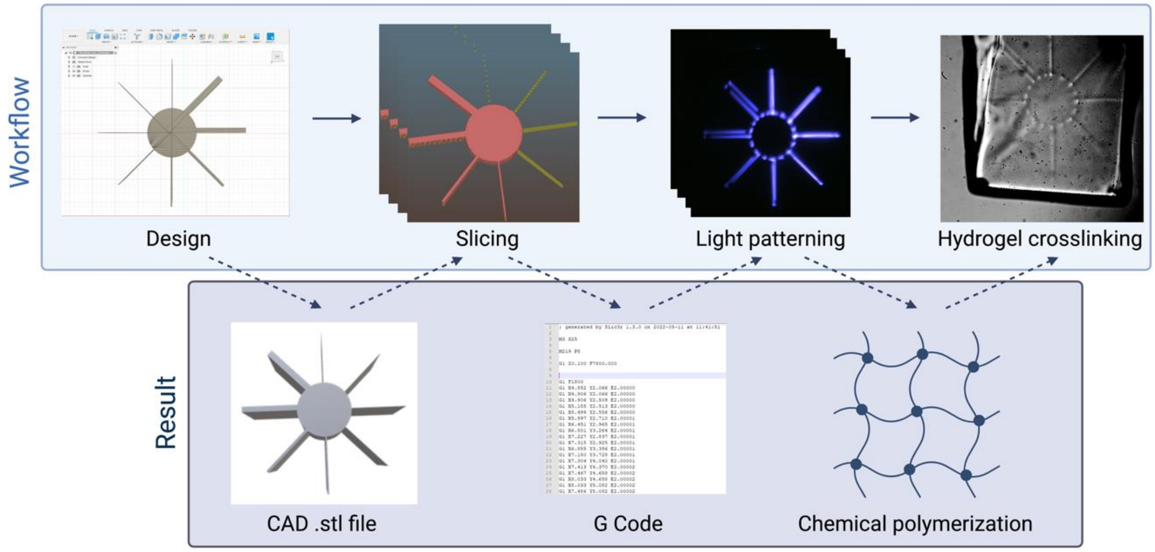

**Fig. S4. The workflow for bioprinting.** It involves four steps from design to crosslinking: designing the 3D object (obtaining a CAD file), slicing the object into z planes (obtaining a G-code), translating into light beam patterning and therefore crosslinking which produces a 3D bioprint. Figure created with Biorender.

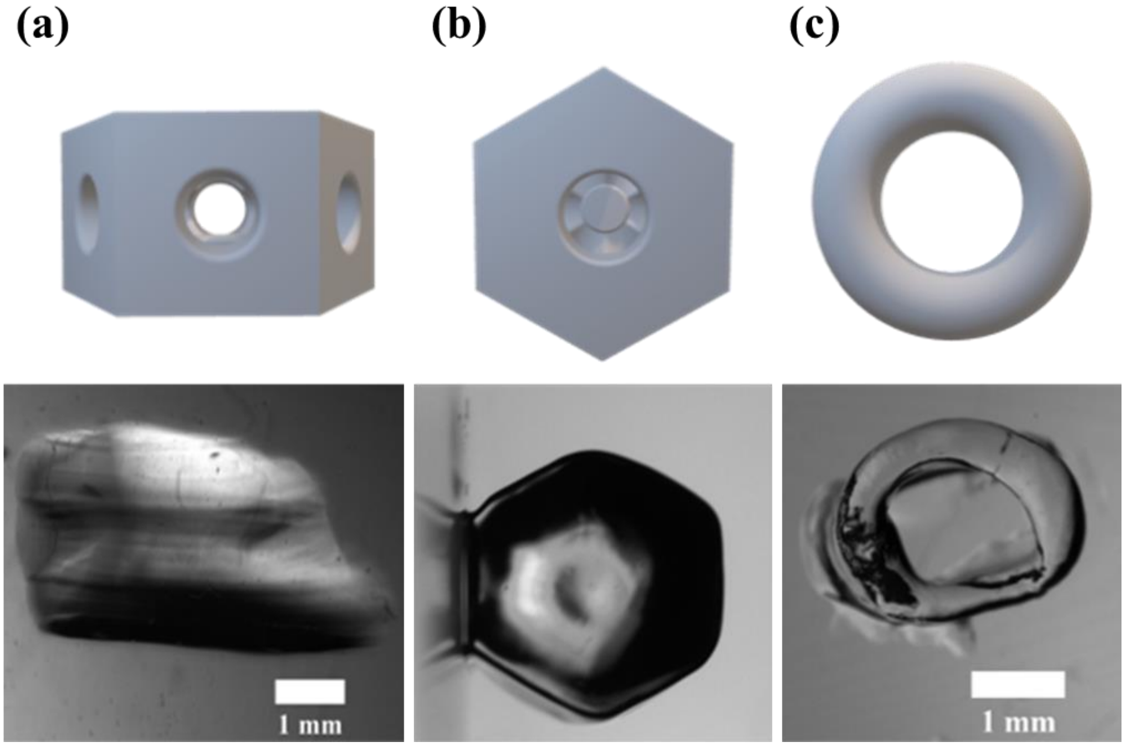

**Fig. S5. Brightfield pictures of the bioprinted constructs.** Constructs were printed using GELMA/PEGDA hydrogel and imaged in air outside of the cuvette. (**a**) Side view and (**b**) top view of the liver lobule. (**c**) Top view of the torus. The resolution wheel did not withstand the extraction process and could not be imaged; however, the liver lobule and torus maintained their shape. Microscope: Zeiss SteREO Discovery V8. Objective: Plan Apo S, 0.63×/0.116 FWD 81 mm. Camera: AxioCam IcC SIN.

(a)

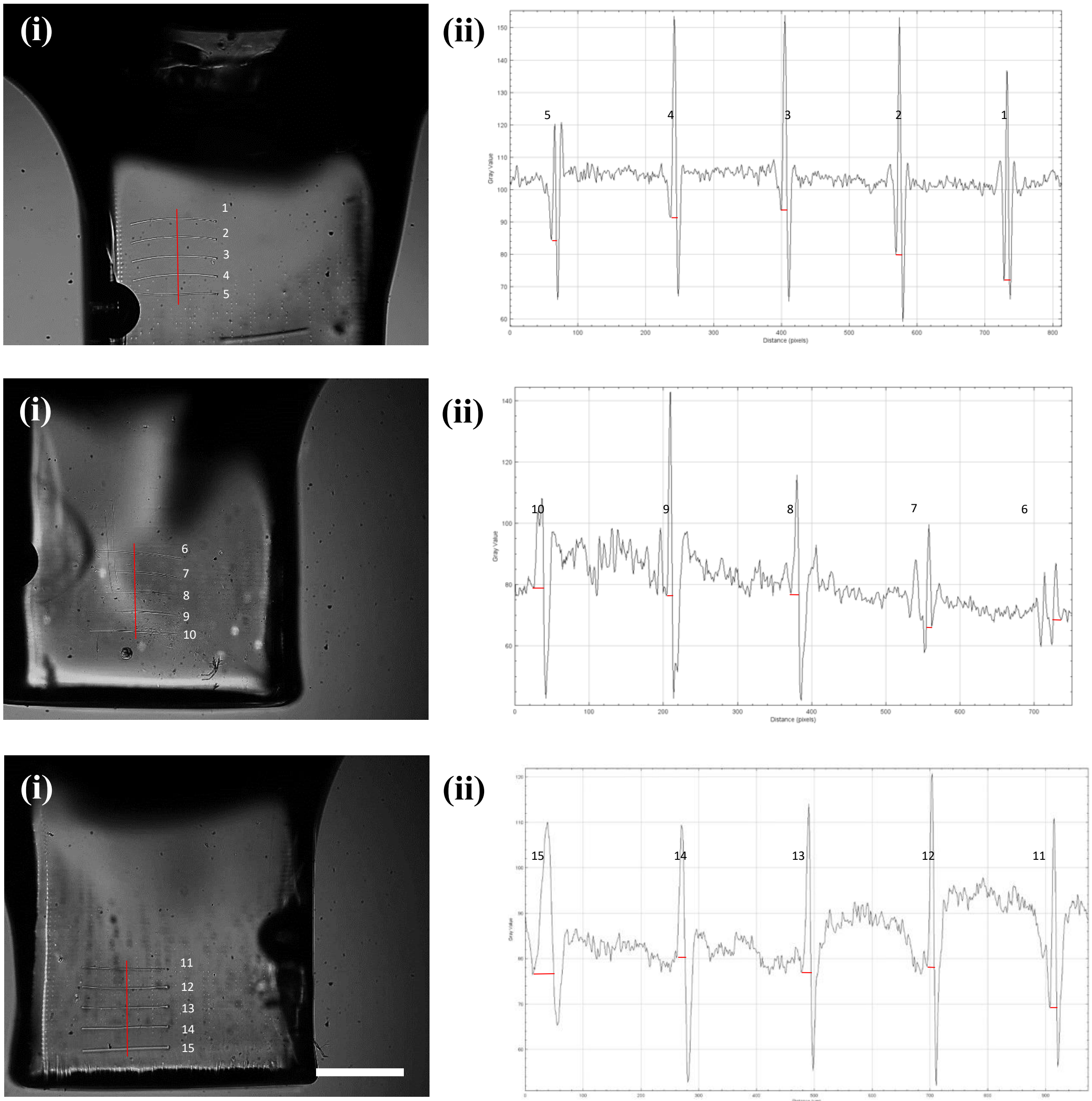

(d)

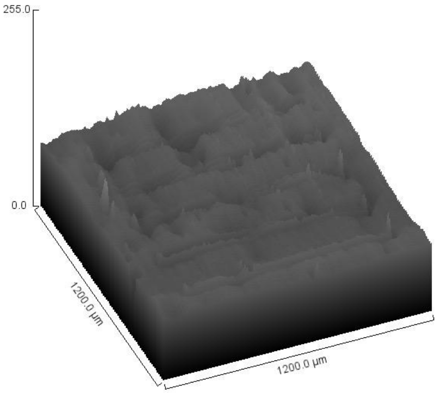

(e)

| | Width [µm] |
|---|---|
| Mean | 15.7 |
| Median | 14.1 |
| St. Dev. | 9.1 |

**Fig. S6. Printing resolution of the patterned light-beam with resin.** (**a**), (**b**) and (**c**) represent five lines printed as triplicates, with (**a**) – (**c**)(**i**) being the brightfield image of the printed lines and (**a**) – (**c**)(**ii**) the analysis of the width of each line. Red lines in (**a**) – (**c**)(**i**) show where the measurement for the width was conducted, red lines in (**a**)(**ii**) – (**c**)(**ii**) show the measured widths for each line in ImageJ and numbers referring to the lines from brightfield images in (**a**)(**i**) – (**c**)(**i**). (**d**) Example surface plot for (**a**)(**i**). (**e**) Calculations for mean, median and standard deviation for the width of the 15 lines printed. Brightfield images captured in the bioprinter. Scale bar: 1 mm.

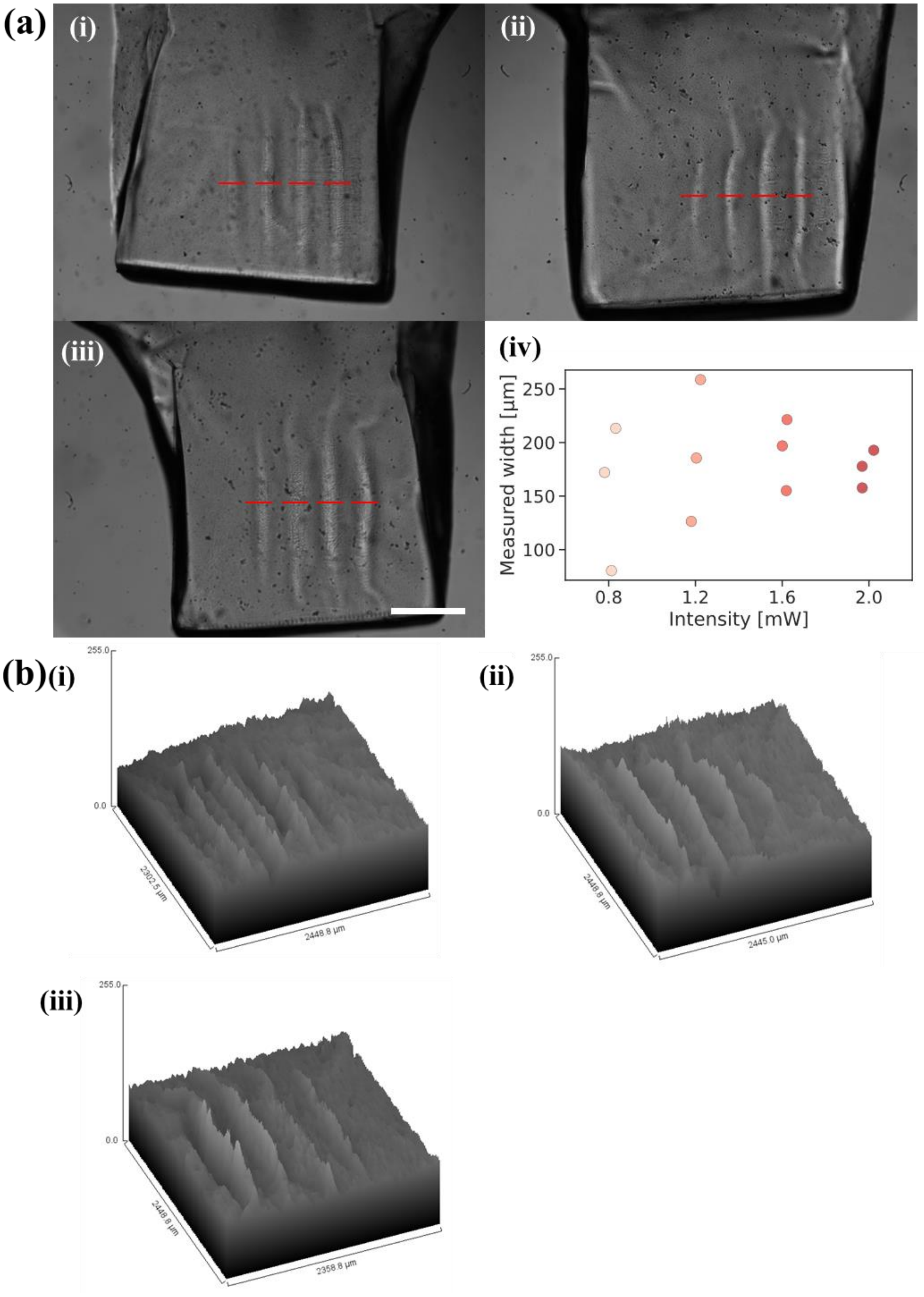
(a)
(i)
(ii)
(iii)
(iv)
Measured width [μm]
250
200
150
100
0.8
1.2
1.6
2.0
Intensity [mW]
(b)
255.0
0.0
2302.5 μm
2448.8 μm
2445.0 μm
2358.8 μm

**Fig. S7. Measurements of the crosslinked orthogonal static light sheet.** It was printed using Cellendes hydrogel 1 with 0.8, 1.2, 1.6 and 2 mW, respectively. (**a**)(**i**), (**ii**) and (**iii**) Three replicates à five laser power settings (2, 1.6, 1.2, 0.8, 0.4 mW) were conducted, respectively. For 0.4 mW no crosslinked sheet was observed. Red lines indicate the position of measurement. (**a**)(**iv**) The total average for the sheet width was measured to be 178.2 µm ± 46.2 µm, with the smallest sheet at 80.8 µm width using 0.8 mW of laser power. (**b**)(**i**), (**ii**) and (**iii**) Surface plots for (**a**)(**i**) to (**iii**). Brightfield images captured in the bioprinter. Scale bar: 1 mm.

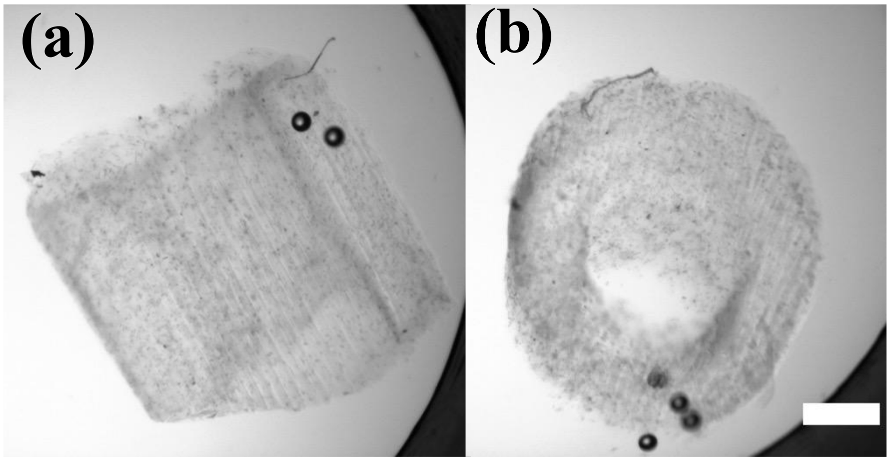

**Fig. S8. Stereomicroscope images of the bioprinted cylinder**. (**a**) Side view of the construct. The bottom of the object was truncated during manipulation. (**b**) Top view of the construct. The hollow part of the cylinder is well resolved and the object is not collapsing. Microscope: Zeiss SteREO Discovery V8. Objective: Plan Apo S, 0.63×/0.116 FWD 81 mm. Camera: AxioCam IcC SIN. Scale bar: 500 µm.

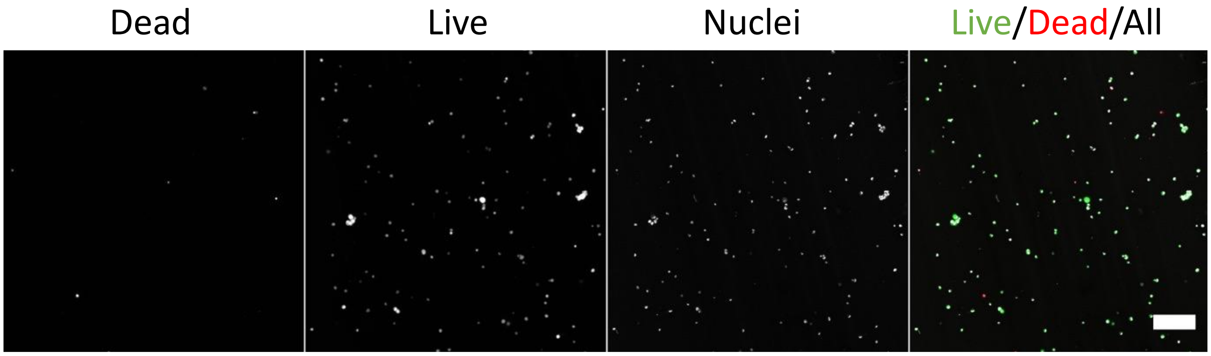

**Fig. S9. Image of the live dead assay of Hs27 fibroblasts encapsulated in Cellendes hydrogel 2 right after bioprinting (day 0).** The dead cells are stained with propidium iodide (PI), the live cells with fluorescein diacetate (FDA) and the nuclei with Hoechst 33342. Microscope: Zeiss AxioObserver LSM780. Objective: Plan ApoChromat 20×/0.8 M27. Voxel size “Hs27 Single Beam”: 0.52 × 0.52 × 6 µm. Voxel size “Hs27 Beam & static LS” and “Co-culture Hs27 & HaCaT”: 0.83 × 0.83 × 6 µm. Scale bar: 100 µm.

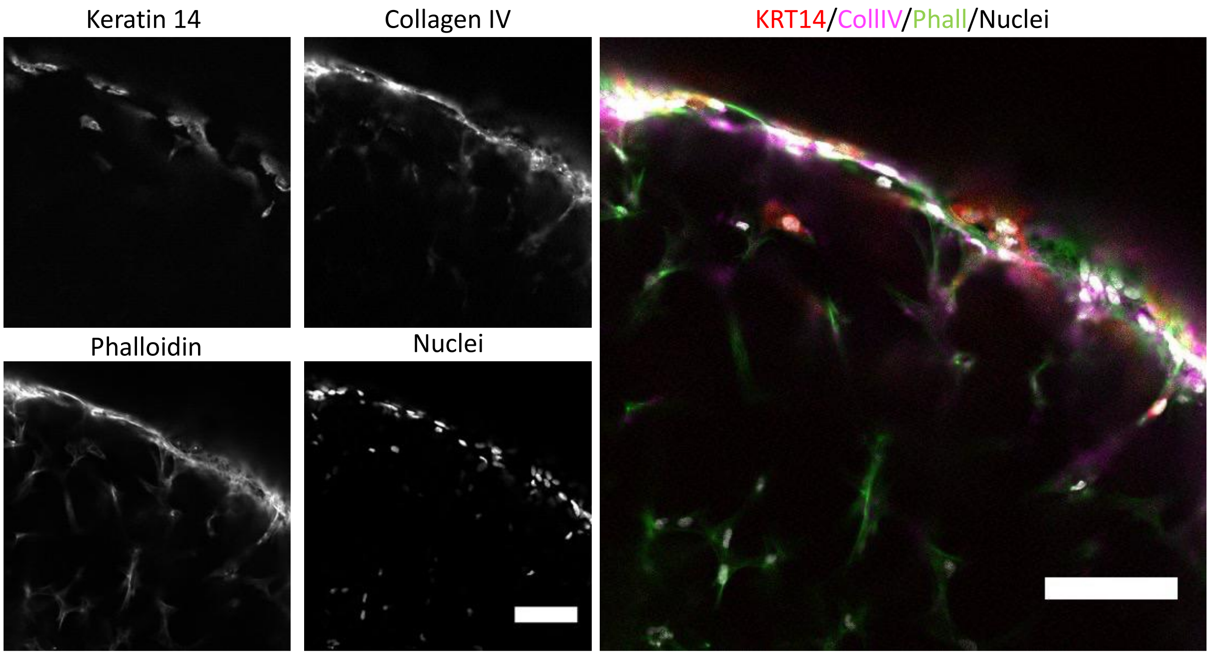

**Fig. S10. Cross section of the cylinder with immunofluorescently stained fibroblasts (Hs27) and keratinocytes (HaCaT).** Keratin 14 (KRT14), Collagen IV (CollIV), phalloidin (Phall) and the nuclei (Hoechst 33342) were stained. A layer of keratinocytes, identifiable by the presence of Keratin 14, is forming on the outside of the object. Microscope: Zeiss AxioObserver LSM780. Objective: Plan ApoChromat 20×/0.8 M27. Scale bar: 100 µm.

**Theoretical principle of Light sheet bioprinting**

The photopolymerization process is limited by the photon density in the print volume. A theoretical photon density threshold can be specified by determining the light sheet properties. Multiple definitions of the beam waist radius $\omega_0$ exist (FWHM, 1/e2, D4σ). The 1/e2 width is a reasonable approximation:

$$\omega_0 \cong (1 - e^{-2}) \times \frac{\lambda_{exc}}{2 \times NA_{ill}}$$

Using the objective lens properties from the setup ($NA_{ill}$ = 0.06) and the excitation wavelength of the laser beam $\lambda_{exc}$ = 395 nm, a theoretical beam waist radius $\omega_0$ of 2846 nm and a beam waist diameter $\varnothing_{beam}$ of 5692 nm was calculated. The Rayleigh length $x_R$ is defined as the distance along the propagation of a gaussian beam from its waist to the spot where the area of its cross section is doubled:

$$x_R = \frac{n \times \pi \times \omega_0^2}{\lambda_{exc}}$$

Inserting the refractive index for water (n = 1.33), the beam waist radius $\omega_0$ (2846 nm) and the excitation wavelength of the laser beam $\lambda_{exc}$ = 395 nm a Rayleigh length of 85690 nm or 85.69 µm was calculated. One light sheet $x_{FOV}$ can be approximated by multiplying the Rayleigh length by two, resulting in 171 µm.

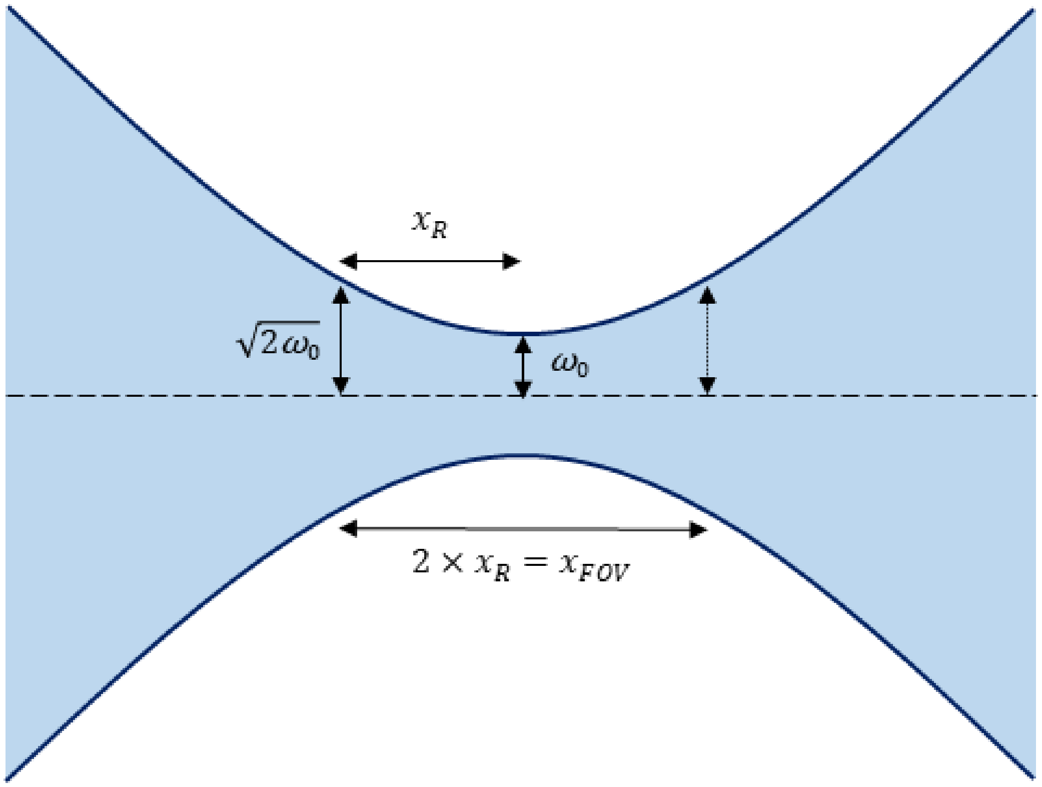

**Fig. S11. Gaussian beam profile**. Beyond the Rayleigh length, the beam cross-section increases almost fourfold with increasing distance from the focal plane. Thus, the number of photons per unit area also decreases in the same order, resulting in a non-linear reduction of radicals and at the same time in a non-linear photo crosslinking process (*69,70*).

**Table S1. Laser power measurements for the single laser beam.**

| S-Value | Power in % | Measured power [mW] | Measured power - Blank [mW] | Designed Power [mW] | Difference: Designed - Measured power [mW] | Percentage of power left at print focal spot |
|---|---|---|---|---|---|---|
| 255 | 100 | 23.27 | 23.27 | 60 | 36.7 | 38.78 |
| 242 | 95 | 22.16 | 22.16 | 57 | 34.8 | 38.87 |
| 230 | 90 | 20.91 | 20.91 | 54 | 33.1 | 38.72 |
| 217 | 85 | 19.95 | 19.95 | 51 | 31.1 | 39.11 |
| 204 | 80 | 18.6 | 18.60 | 48 | 29.4 | 38.75 |
| 191 | 75 | 17.62 | 17.62 | 45 | 27.4 | 39.15 |
| 179 | 70 | 16.33 | 16.33 | 42 | 25.7 | 38.88 |
| 166 | 65 | 15.2 | 15.20 | 39 | 23.8 | 38.97 |
| 153 | 60 | 14.13 | 14.13 | 36 | 21.9 | 39.25 |
| 140 | 55 | 12.9 | 12.90 | 33 | 20.1 | 39.09 |
| 128 | 50 | 11.85 | 11.85 | 30 | 18.2 | 39.50 |
| 115 | 45 | 10.56 | 10.56 | 27 | 16.4 | 39.11 |
| 102 | 40 | 9.49 | 9.49 | 24 | 14.5 | 39.54 |
| 89 | 35 | 8.23 | 8.23 | 21 | 12.8 | 39.18 |
| 77 | 30 | 7.18 | 7.18 | 18 | 10.8 | 39.88 |
| 64 | 25 | 5.89 | 5.89 | 15 | 9.1 | 39.26 |
| 51 | 20 | 4.81 | 4.81 | 12 | 7.2 | 40.07 |
| 38 | 15 | 3.62 | 3.62 | 9 | 5.4 | 40.21 |
| 25 | 10 | 2.47 | 2.47 | 6 | 3.5 | 41.14 |
| 13 | 5 | 1.27 | 1.27 | 3 | 1.7 | 42.28 |
| 0 | 0 | 0.0015 | 0 | 0 | 0.0 | - |

**Table S2. Laser power measurements for the static light sheet beam.**

| S-Value | Power in % | Measured power [mW] | Measured power - Blank [mW] | Designed Power [mW] | Difference: Designed - Measured power [mW] | Percentage of power left at print focal spot |
|---|---|---|---|---|---|---|
| 51.0 | 20 | 4.240 | 4.239 | 10 | 5.8 | 42.39 |
| 38.3 | 15 | 3.180 | 3.179 | 7.5 | 4.3 | 42.38 |
| 25.5 | 10 | 2.110 | 2.109 | 5 | 2.9 | 42.17 |
| 12.8 | 5 | 1.045 | 1.044 | 2.5 | 1.5 | 41.74 |
| 2.6 | 1 | 0.181 | 0.180 | 0.5 | 0.3 | 35.90 |
| 2.3 | 0.9 | 0.162 | 0.161 | 0.45 | 0.3 | 35.67 |
| 2.0 | 0.8 | 0.141 | 0.140 | 0.4 | 0.3 | 34.88 |
| 1.8 | 0.7 | 0.117 | 0.116 | 0.35 | 0.2 | 33.00 |
| 1.5 | 0.6 | 0.101 | 0.100 | 0.3 | 0.2 | 33.17 |
| 1.3 | 0.5 | 0.083 | 0.082 | 0.25 | 0.2 | 32.60 |
| 1.0 | 0.4 | 0.062 | 0.061 | 0.2 | 0.1 | 30.25 |
| 0.8 | 0.3 | 0.046 | 0.045 | 0.15 | 0.1 | 29.67 |
| 0.5 | 0.2 | 0.032 | 0.031 | 0.1 | 0.1 | 30.50 |
| 0.3 | 0.1 | 0.020 | 0.019 | 0.05 | 0.0 | 37.00 |

**Table S3. Energy calculations for every 3D print in this publication.**

| Printed object | S-Value | Power in % | Measured power [mW] | Energy [mJ/cm$^2$] | Exposure time [s] |
|---|---|---|---|---|---|
| Wheel of resolution (Hydrogel) | 45 | 17.6 | 4.22 | 5.02 | - |
| Liver lobule (Hydrogel) | 51 | 20 | 4.81 | 5.72 | - |
| Torus (Hydrogel) | 60 | 23.5 | 5.59 | 6.65 | - |
| Hollow cylinder (Hydrogel resolution)<br>(hydrogel with cells – medium pellet<br>– big pellet) | 45<br>51<br>92 | 17.6<br>20<br>36 | 4.22<br>4.65<br>8.37 | 5.02<br>5.72<br>10.30 | -<br>-<br>- |
| Five lines (Resin) | 45 | 17.6 | 4.22 | 5.02 | - |
| Static LS (Resin) | - | 0.1 | 0.02 | 0.017; 0.013; 0.010; 0.007; 0.004 | 5; 4; 3; 2; 1 |
| Static LS (Hydrogel) | 51; 41; 31; 20; 10 | 20; 16; 12; 8; 4 | 4.24; 3.39; 2.53; 1.68; 0.83 | 7.35; 6.88; 4.38; 2.91; 1.44 | 10 |

**Table S4. List of antibodies and dyes**

| Component | Supplier | Stock | Dilution |
|---|---|---|---|
| MitoTracker red FM | Invitrogen | 1 mM | 1:1000 |
| Fluorescein diacetate | Sigma-Aldrich Chemie GmbH | 5mg/ml in acetone | 1:500 |
| Propidium iodide | Sigma-Aldrich Chemie GmbH | 1 mg/ml in PBS | 1:100 |
| Hoechst 33342 | ThermoFisher Scientific | - | 1:500 |
| Alexa Fluor 546 Phalloidin | ThermoFisher Scientific | - | 1:200 |
| Alexa Fluor 647 Phalloidin | ThermoFisher Scientific | - | 1:100 |
| Collagen IV (rabbit) | Abcam | 1 mg/ml | 1:400 |
| Cytokeratin 14 (mouse) | Santa Cruz Technologies | 200 µg/ml | 1:50 |
| Vimentin (rabbit) | Cell Signaling | - | 1:100 |
| Anti-rabbit Alexa Fluor 488 (goat) | ThermoFisher Scientific | 2 mg/ml | 1:400 |
| Anti-mouse Alexa Fluor 568 (donkey) | ThermoFisher Scientific | 2 mg/ml | 1:400 |

**Table S5. Composition of the hydrogel used for resolution experiment (Cellendes hydrogel 1)**

| Gel components | Final concentration |
|---|---|
| RGD-N-Dex | 3.5 mM (reactive group) |
| CD-HyLink | 5 mM (reactive group) |
| LAP | 0.5 mM |
| Agarose | 0% |

**Table S6. Composition of the hydrogel used for resolution experiment (GelMA/PEGDA)**

| Gel components | Final concentration |
|---|---|
| GelMA | 10%(w/v) |
| PEGDA | 10%(w/v) |
| LAP | 0.2%(w/v) |
| Tartrazine | 0.0125%(w/v) |

**Table S7. Composition of the hydrogel used for hydrogel testing and cell encapsulation (Cellendes hydrogel 2)**

| Gel components | Final concentration |
|---|---|
| RGD-N-Dex | 3.5 mM (reactive group) |
| CD-HyLink | 3.5 mM (reactive group) |
| LAP | 0.5 mM |
| Agarose | 0.1% |

**Table S8. Comparison of the BRIGHTER light sheet bioprinter with other state of the art 3D (bio-)printers.**

| 3D (Bio)printer | Technique | Speed [$mm^3/s$] | Print time increase with volume | Resolution [µm] | Energy Dose [$mJ/cm^2$] | Cell viability [%] | Print time for 100 $mm^3$ with best resolution [s] | Print time for 100 $mm^3$ with best resolution [min] | Reference |
|---|---|---|---|---|---|---|---|---|---|
| LS-Bioprinter | orthogonal laser-patterning | 0.66 | Linear | 15.7 | 5 - 10 | >90 | 152 | 2.53 | This work |
| Extrusion | Fused deposition modelling | 10 - 20 | Linear | >100 | - | >70 | 7 | 0.11 | Gómez-Blanco et al. 2021 |
| Stereolithographic (SLA) | LCD photomask | 0.21 | Linear in height | 100 - 200 | 50 | >95 | 476 | 7.9 | Breidebandet al. 2022 |
| Stereolithographic (SLA) | Digital Light processing (DLP) | 0.018 | Linear in height | 25 | 10 - 100 | >85 | 5556 | 92.6 | Torras et al. 2022 |
| Two Photon | Two Photon-Photopolymerization | 0.000045 - 0.125 | Linear | 0.8 - 10 | NA | >70 | 4444444 | 74074 | Dobos et al. 2020 |
| Volumetric Bioprinting | Tomographic additive manufacturing | 6 - 182 | Same time up to 3.9 $cm^3$ (14 x 14 x 20 mm) | 40 - 100 | 100 - 500 | >90 | 22.7 | 0.38 | Loterie et al. 2020, Bernal et al. 2022, Gehlen et al. 2023 |
| Volumetric Bioprinting | Xolography | 55 | Same time up to 1 $cm^3$ | 20 | 50 - 300 | Not suitable for bioprinting | 2 | 0.03 | Regehly et al. 2020 |

**Movie S1. Video of the patterning of a resolution wheel, as seen from the integrated camera (view from the side, patterning not at full speed).**

**Movie S2. Static light sheet and laser patterning beam (view from the top, patterning not at full speed).**

**Movie S3. Z-stack of the Hs27-mitotracker and HUVEC-GFP co-culture imaged in the light sheet part of the bioprinter (color-coded by depth). Only the 488 nm channel showing the HUVEC-GFP cells is here shown. Voxel size: 0.69 × 0.69 × 10 µm. Objective lenses: Zeiss A-Plan 2.5x/0.06 (excitation).**
